# Paralog protein compensation preserves protein-protein interaction networks following gene loss in cancer

**DOI:** 10.1101/2024.09.26.615228

**Authors:** Anjan Venkatesh, Niall Quinn, Swathi Ramachandra Upadhya, Barbara De Kegel, Alfonso Bolado Carrancio, Thomas Lefeivre, Olivier Dennler, Kieran Wynne, Alexander von Kriegsheim, Colm J. Ryan

## Abstract

Proteins operate within dense interconnected networks, where interactions are necessary both for stabilising proteins and for enabling them to execute their molecular functions. Remarkably, protein-protein interaction networks operating within tumour cells continue to function despite widespread genetic perturbations. Previous work has demonstrated that tumour cells tolerate perturbations of paralogs better than perturbations of singleton genes, but the mechanisms behind this genetic robustness remains poorly understood. Here, we systematically profile the proteomic response of tumours and tumour cell lines to gene loss. We find many examples of active compensation, where deletion of one paralog results in increased abundance of another, and collateral loss, where deletion of one paralog results in reduced abundance of another. Compensation is enriched among sequence-similar paralog pairs that are central in the protein-protein interaction network and widely conserved across evolution. Compensation is also significantly more likely to be observed for gene pairs with a known synthetic lethal relationship. Our results support a model whereby loss of one gene results in increased protein abundance of its paralog, stabilising the protein-protein interaction network. Consequently, tumour cells may become dependent on the paralog for survival, creating potentially targetable vulnerabilities.

## Introduction

Tumour cells tolerate enormous amounts of genetic perturbations, including loss of function mutations and deletions of protein-coding genes^1–4^. We, and others, have demonstrated that duplicate genes (paralogs) contribute significantly to this genetic robustness – tumour cells are more tolerant to loss of paralogs than singleton genes, resulting in deletions of paralogs being more frequently observed in tumour genomes^4,5^. Perhaps the simplest explanation for the increased dispensability of paralogs is that pairs of paralogs can compensate for each other’s loss because of their shared functions. Direct evidence for this model comes from double perturbation screens in cancer cell lines – often either paralog in a pair can be lost individually with relatively little fitness consequence, but their combined inhibition causes cell death, a phenomenon termed synthetic lethality^6–10^. The increased dispensability of paralog genes, combined with evidence of frequent synthetic lethality between paralog pairs, supports a model whereby tumours can tolerate the loss of an individual paralog because its counterpart can compensate for its loss. This provides a clear model for the tolerance of tumour cells for genetic perturbation but provides no insight into the mechanisms by which this compensation takes place. In particular, it is entirely unclear how the molecular interaction networks operating within tumour cells adapt to the loss of paralog genes. Is there a need for increased production of the compensating paralog in order for it to carry out the functions of the lost paralog? How do protein complexes, which often rely on balance between different subunits, adapt to the loss of paralogous subunits?

To our knowledge, no systematic effort to understand the mechanisms behind paralog compensation has been undertaken in the context of cancer or human cells. However, there have been a number of systematic efforts to do so in the budding yeast *Saccharomyces cerevisiae*. Early work identified that some pairs of paralogs exhibit *transcriptional reprogramming* such that loss of one paralog was associated with increased transcription of another ^11^. A subsequent systematic study in yeast assessed the proteomic response of paralogs to deletion of their counterpart and found that, of 202 paralog pairs tested, just over 10% exhibit ‘*need based upregulation’* such that deletion of one paralog caused increased protein abundance of the other^12^. Intriguingly, the pairs that exhibited this *upregulation* were enriched for pairs known to be synthetic lethal, suggesting a relationship between phenotypic compensation and molecular compensation. This study also identified a smaller number of examples of the opposite effect, where deletion of one gene was associated with reduced protein abundance of its paralog. The authors speculated that these relationships may be due to stoichiometric requirements of protein complexes containing both paralogs. A subsequent systematic effort to understand how loss of one paralog alters the protein-protein interactions of its counterpart revealed examples of both compensation and the opposite effect, i.e. dependency, with the two relationships occurring at similar frequencies ^13^. In the case of compensation, loss of one paralog resulted in increased interaction between the compensatory paralog and the protein-protein interaction partners of the lost paralog. In the case of dependency, deletion of one paralog resulted in the other paralog losing protein-protein interaction partners. Further analysis suggested that pairs that exhibit dependency are enriched among paralogs that form heteromers and that these pairs may stabilise each other through this interaction^14^.

While no systematic study of the molecular consequences of paralog loss has been performed in cancer, there have been systematic efforts to understand how tumour cells respond to copy number variation in general^15–17^. These studies have made use of matched genomic, transcriptomic, and proteomic profiles of tumours to understand how genetic variation influences transcriptomic and proteomic variation. A particularly striking finding has been that copy number changes at the genomic level are often attenuated at the protein level and that this is especially evident for genes that encode protein complex subunits, suggesting that regulatory mechanisms operate within tumour cells to maintain protein complex stoichiometry^15,16,18,19^. An additional finding has been that many protein-protein interaction partners control each other’s abundance, such that loss of one gene was associated with reduction of another^15^. There is some evidence that the regulation of protein complex stoichiometry and the controlling relationships between paralog pairs occurs via proteasome-mediated degradation of excess subunits^15,16,19^.

Here we sought to directly address the question of how tumour cells respond to loss of paralog genes. First, we investigated the proteomic consequences of knocking out a set of 34 paralogous genes using CRISPR-Cas9 gene editing in a single genetic background. Consistent with results from yeast, we identified examples of both collateral loss and protein compensation. We then performed a larger scale analysis of tumour samples with matched genomes, transcriptomics, and proteomes. This allowed us to assess the regulatory relationships between thousands of paralog pairs and revealed hundreds of examples of collateral loss and protein compensation. The set of compensation pairs was enriched in paralogs that were highly conserved across evolution, central in the protein-protein interaction network, and were members of essential protein complexes. In contrast, paralogs that exhibit collateral loss relationships were less conserved, more peripheral on the protein-protein interaction network, and less likely to be involved in essential functions. Compensation pairs were also highly enriched among known synthetic lethal pairs. Overall, our results suggest a model where gene loss-induced disruptions to the protein-protein interaction network are buffered by increased protein abundance of paralogous proteins, upon which tumour cells then become dependent for survival. These proteomic compensation events are widespread in cancer, enabling cancer cells to thrive despite frequent gene loss.

## Results

### Mass spectrometry profiling of isogenic knockouts reveals compensation and collateral loss events

We first modelled the proteomic consequences of paralog loss using isogenic cell lines, allowing us to causally link an introduced mutation to changes in protein abundance. Previous systematic efforts to identify associations between the loss of one protein and the abundance of a paralog had been carried out in budding yeast and were performed by mating GFP-tagged strains (where fluorescence was used as a proxy for protein abundance) with gene deletion strains^12^. Since this approach does not readily translate to mammalian cells, we used an alternative approach – CRISPR perturbation of individual paralogs followed by unbiased quantification of the proteome by mass spectrometry. We used the HAP1 model for this work – HAP1 is a near-haploid cell line, allowing for reliable generation of non-heterozygous mutations. We selected 34 paralogous genes to mutate, based on a number of criteria – firstly the genes had to be detected in the HAP1 transcriptome but be non-essential for growth^20^. Within this subset, we prioritised genes that had a previously reported or suspected synthetic lethal interaction with at least one of their paralogs, on the grounds that these pairs exhibit phenotypic compensation and therefore might be more likely to display proteomic compensation^6–10,12,21^. We performed whole-cell lysate proteomic profiling of 34 HAP1-derived clones, each of which had a frameshift insertion or deletion of a single paralog, and the parental HAP1 strains for comparison(Fig. 1a, Methods). On average, we quantified 4,544 proteins per clone profiled (Supplementary Table 1). We first verified that the mutation in each of the 34 profiled clones caused a reduction in the abundance of the associated protein. For 23 (∼68%) of the models, we were able to verify a significant drop in the abundance of the protein product of the targeted gene (Supplementary Table 2). For the remainder we could either not reliably quantify the protein in the parental HAP1 model (8/34 models), so a reduction in the protein abundance could not be assessed, or we could quantify the protein but not verify an associated drop in abundance (3/34 models).

**Figure 1.**
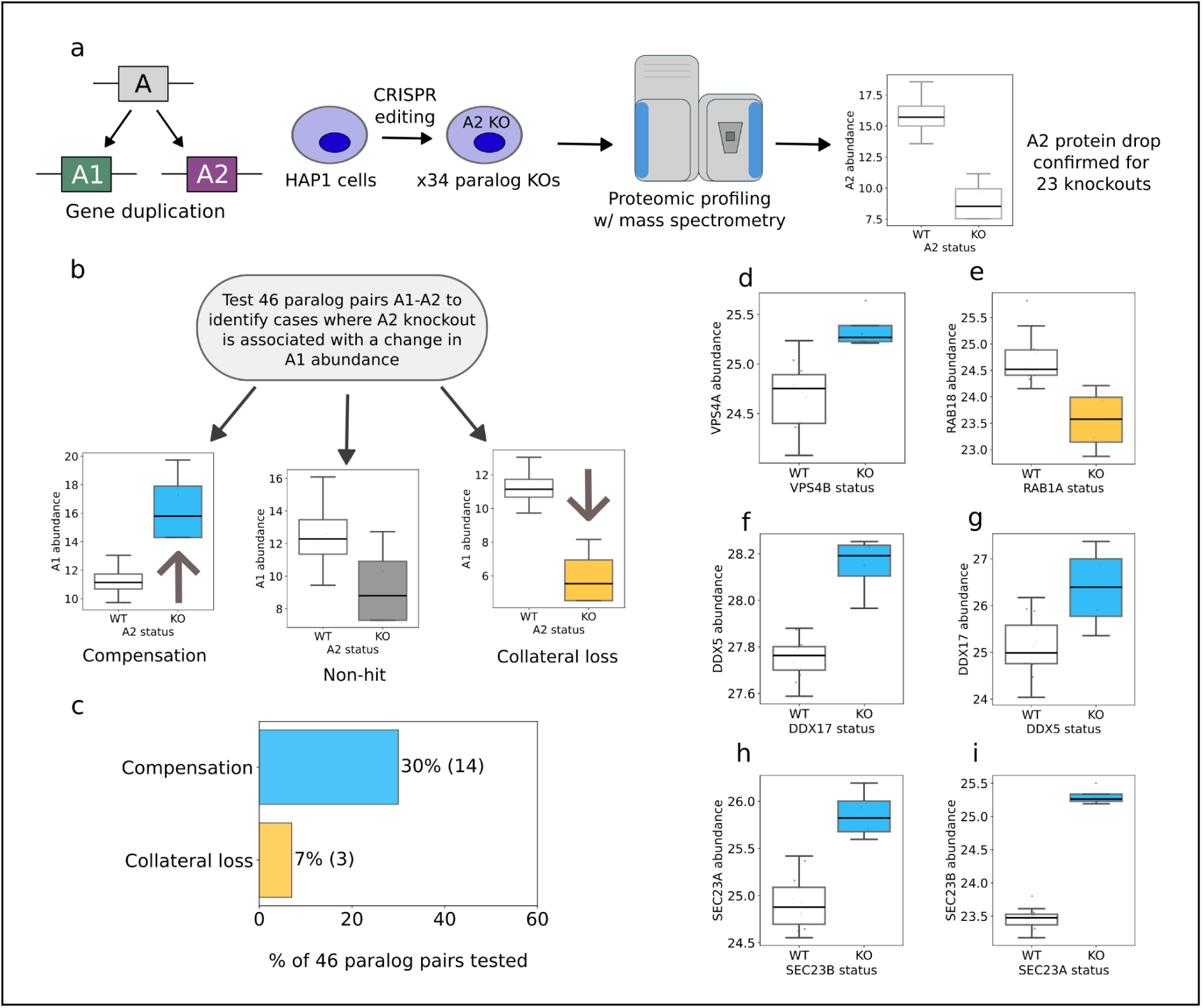
Identifying paralog protein compensation and collateral loss in an isogenic model. **(a)** Experimental setup for proteomic profiling of HAP1 knockouts. **(b)** Workflow for identifying cases of paralog compensation and collateral loss. **(c)** Proportion of compensation and collateral loss pairs identified, with actual count in brackets. **(d-i)** Examples of compensation and collateral loss hits showing increase (blue) or decrease (yellow) in the abundance of a protein following the knockout of its paralog.

For the 23 genes whose loss-of-function mutation was proteomically verified, we then tested whether the mutation caused a change in the abundance of each of their detected paralogs (see Methods). In total we tested 46 gene pairs, since many of the targeted genes had multiple identifiable paralogs (Fig. 1b). Using this approach, we identified 14 gene pairs with compensation relationships and 3 gene pairs with collateral loss relationships (Fig. 1c, Supplementary Table 3). Here, compensation refers to a pair *A1*-*A2* where a protein A1 has significantly (FDR < 5%; p < 0.05) increased abundance when its paralog *A2* is knocked out, and collateral loss refers to a pair where A1 has significantly reduced abundance when *A2* is knocked out. For example, we identified a compensatory relationship between *VPS4B* and VPS4B (which encode ATPases involved in ESCRT-dependent membrane remodelling), a pair we and others have previously identified as synthetic lethal^22–24^ (Fig. 1d). In contrast we found that loss of the Rab GTPase *RAB1A* was associated with collateral loss of its paralog Rab GTPase RAB18 (Fig. 1e). In 3 cases where we had knockouts of both genes in a pair, we observed reciprocal compensation effects. For example, in the case of the DEAD-box helicases *DDX5* and *DDX17,* which we have previously identified as synthetic lethal^23^, *DDX17* loss resulted in increased protein abundance of DDX5 (Fig. 1f), while *DDX5* loss resulted in increased protein abundance of DDX17 (Fig. 1g). Similar reciprocal patterns were observed for SEC23A-SEC23B (Fig. 1h, 1i) and SMARCC1-SMARCC2 (Supplementary Table 3).

### Unbiased identification of paralog compensation and collateral loss in a panel of tumour proteomes

Our analysis of HAP1 paralog knockouts demonstrates that both compensation and collateral loss relationships can be detected by mass spectrometry profiling in isogenic models. However, there are some limitations to this approach – most notably, each gene of interest requires the creation of a new isogenic knockout. Due to this lack of scalability, we chose paralog pairs for which we anticipated seeing compensatory effects and our results do not represent an unbiased analysis. To perform a more systematic and unbiased analysis, we next analysed the consequences of paralog loss in a large compendium of tumour proteomes from the Clinical Proteomic Tumor Analysis Consortium (CPTAC). We analysed a set of 930 normalised tumour proteomes with matched transcriptomes and copy number profiles from this project (see Methods). Since these samples come from 8 different studies, each pertaining to a different cancer type, we incorporated cancer type as a covariate when assessing the impact of gene loss on protein abundance. As recurrent homozygous gene loss is relatively uncommon, even in tumours, we focused our analysis on single copy loss^4,25^. For all paralogs that were subject to single copy loss in at least 20 samples, we assessed whether this loss was associated with a reduction in the abundance of the cognate protein. For 2,249 out of 3,116 (∼67%) of tested genes, we found this to be the case (Supplementary Fig. 1, Supplementary Table 4). We next asked, for each of these genes, how the copy number loss impacted the abundance of each of their quantified paralogs (Fig. 2a). In total, we tested 4,568 paralog associations (Supplementary Fig. 2, Supplementary Table 5). At an FDR of 5%, we identified 568 (12%) compensation and 449 (10%) collateral loss hits (Fig. 2b, 2c). For example, loss of the DNA topoisomerase *TOP2B* was associated with a significant increase of TOP2A protein (Fig. 2d, Supplementary Fig. 1b)^26^. In contrast, loss of the GDP-mannose pyrophosphorylase *GMPPB* was associated with a significant decrease in the abundance of its paralog GMPPA (Fig. 2g, Supplementary Fig. 1e). GMPPA and GMPPB form a heterodimer to enable GDP-Man synthesis and it is therefore possible that loss of *GMPPB* results in an overall reduction in the abundance of this complex leading to reduced stability of GMPPA^27,28^.

**Figure 2.**
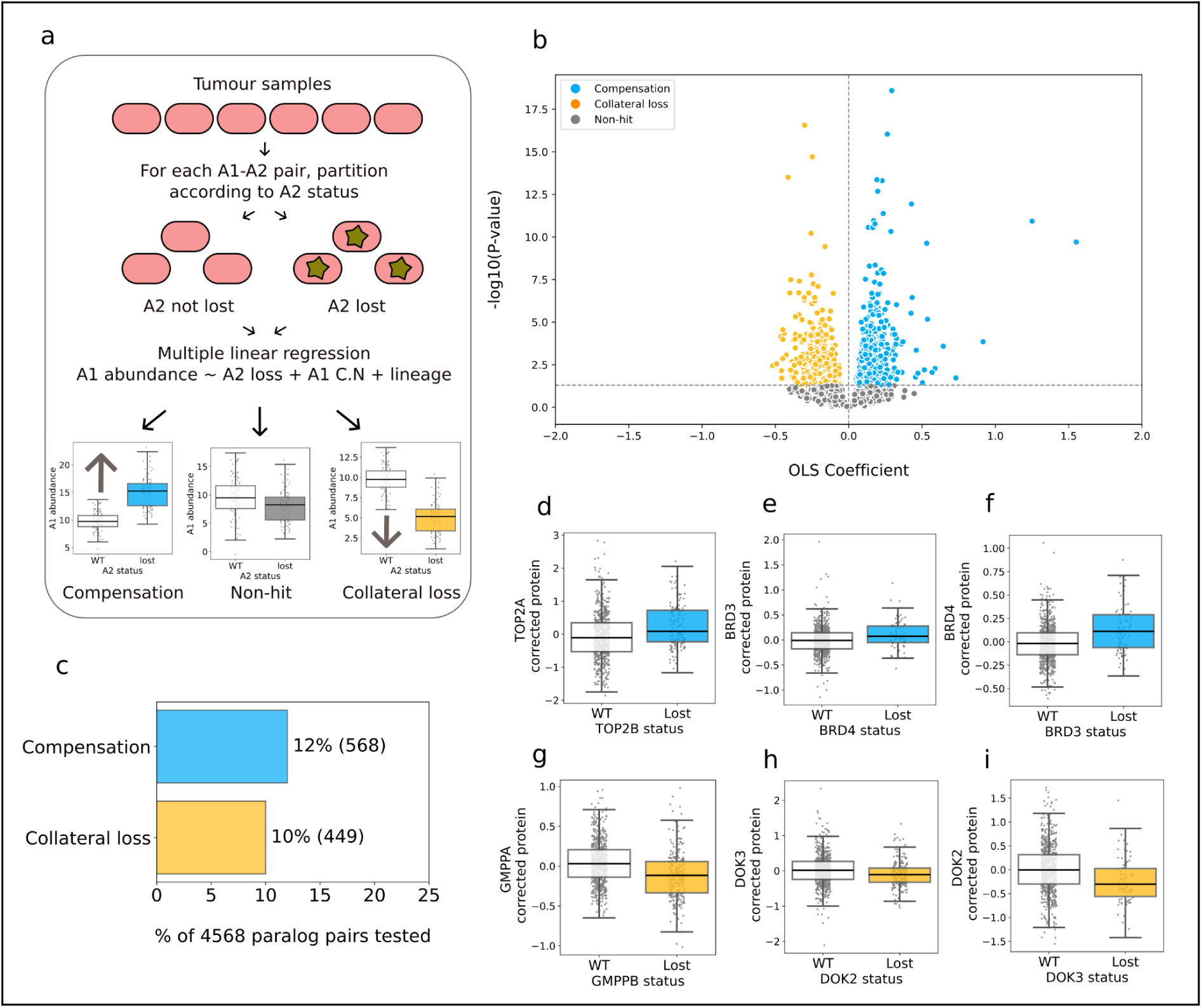
Identifying paralog protein compensation and collateral loss using tumour profiles. **(a)** Workflow diagram for unbiased identification of collateral loss and compensation from tumour proteomic profiles. **(b)** Volcano plot showing -log10(FDR) vs ordinary least squares (OLS) coefficient for all paralog tests run with the CPTAC dataset. Each dot represents a paralog pair. **(c)** Proportion of compensation and collateral loss hits in the CPTAC analysis, with actual counts in brackets. **(d-i)** Boxplots of compensation and collateral loss hits. In each box plot the top and bottom of the box represents the third and first quartiles and the box band represents the median; whiskers extend to 1.5 times the interquartile distance from the box. Each grey dot represents a tumour sample. Position on the y-axis indicates protein abundance adjusted for copy number of the encoding gene and cancer type, which are used as covariates in the regression model.

In our analysis of the HAP1 data, we found that some pairs were observed as compensatory in a symmetric fashion – *A2* loss was associated with an increased abundance of A1, while loss of *A1* was associated with increased abundance of A2. With the CPTAC analysis, most paralog pairs were tested in an asymmetric fashion – i.e. we assessed the impact of *A2* loss on A1’s protein abundance but not *A1*’s loss on A2’s protein abundance. However, we found that, for the subset of pairs that were tested in both directions, there was a significant overlap among reciprocal compensation pairs (OR = 2.9, p = 9e-06; Supplementary Fig. 3a) and a non-significant overlap among collateral loss pairs (OR = 1.6, p = 0.09; Supplementary Fig. 3b). For example, loss of *BRD4* was associated with increased abundance of BRD3 and vice-versa, suggesting reciprocal compensation (Figs. 2e, 2f, Supplementary Figs. 1c, 1d).

In contrast, loss of *DOK3* was associated with reduced protein abundance of DOK2 and vice-versa, suggesting reciprocal collateral loss (Figs. 2h, 2i, Supplementary Figs. 1f, 1g). 3 compensation pairs identified with the unbiased CPTAC analysis were also compensation hits in the HAP1 analysis— SMARCC1-SMARCC2 (i.e. SMARCC1 abundance increases when *SMARCC2* is lost), SEC23B-SEC23A, and DDX5-DDX17 (Supplementary Fig. 4, Supplementary Table 5).

### Compensation pairs are more sequence similar and come from smaller paralog families

We wished to understand what features might be associated with proteomic compensation and collateral loss in the CPTAC dataset. Paralog relationships are typically identified from sequence comparisons and we therefore initially focussed on sequence-derived features. The sequence identity between paralogs is often used as a proxy for functional similarity, with highly sequence-similar proteins presumed to be more functionally similar. We hypothesised that compensatory paralogs might be more functionally similar and therefore display higher sequence identity. We found that this is indeed the case – compensation hits had higher amino acid sequence identity than non-hits (p = 0.02, two-tailed t-test) (Fig. 3a).

**Fig 3.**
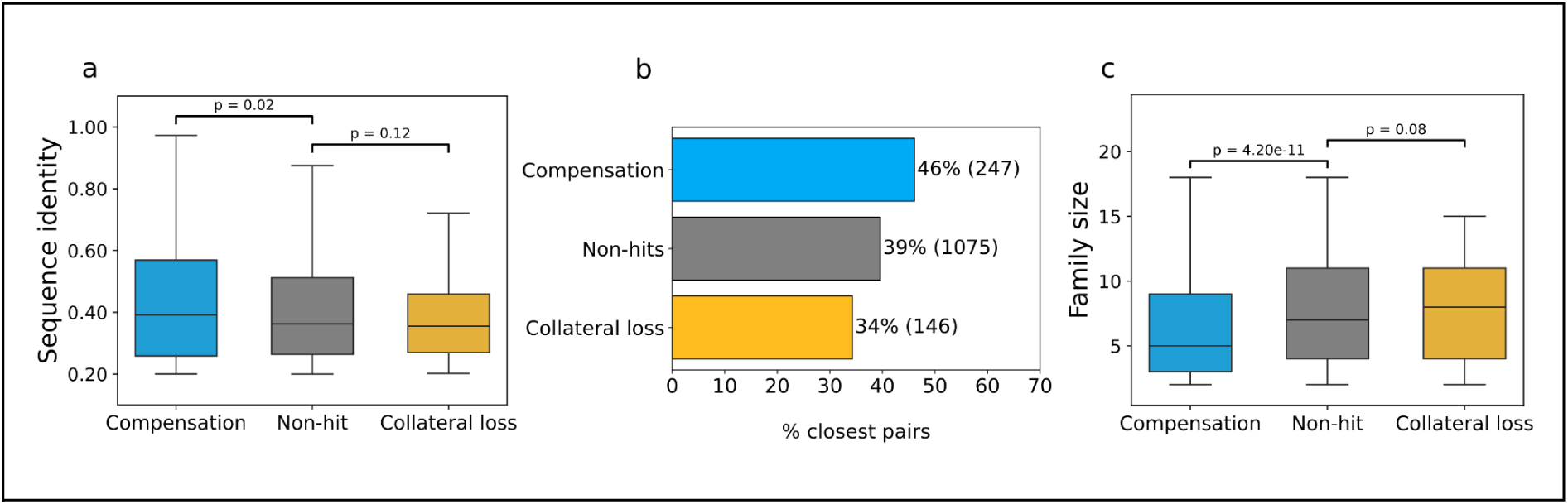
Compensation pairs are more sequence similar, belong to smaller families, and are enriched among closest pairs. **(a)** Box plots showing the amino acid sequence identity of compensation pairs, collateral loss pairs, and non-hits. **(b)** Barchart showing the percentage of paralog pairs in each group that are each other’s ‘closest paralog’ among all members of a paralog family. All p-values are from two-t t-tests (assuming equal variances). **(c)** Boxplots showing the family size (number of paralogs belonging to the same family) for pairs with compensation, collateral loss, and non-hits.

Due to multiple rounds of duplication, often a single gene can have multiple paralogs. We anticipated that compensation might be more common among ‘closest pairs’ – i.e. pairs which are each other’s most sequence-similar paralog within a larger gene family. We found that this is indeed the case for compensation pairs (Fisher’s Exact Test, OR = 1.4; p = 1.2e-03) (Fig. 3b), while collateral loss pairs are less likely to be closest pairs (OR = 0.8, p = 0.045). Furthermore, we found that compensation was enriched among smaller paralog families (t-test p = 4.2e-11) (Fig. 3c).

### Compensation pairs are more central in the protein-protein interaction network and perform essential functions

We hypothesised that proteomic compensation might be a means of stabilising the protein-protein interaction network against perturbations. If this is the case, one might anticipate that proteins with more interactions should be more likely to display compensatory relationships. We therefore calculated the degree centrality (total interactions normalised by network size) of all paralogs using protein-protein interactions from the BioGRID^29^. We found that the genes lost in compensation pairs (i.e. compensated-for genes) tend to have significantly higher protein-protein interaction degree centrality (t-test p = 1.6e-08) (Fig. 4a). In contrast, genes associated with collateral loss relationships had significantly lower degree centrality (t-test p = 1.2e-04) (Fig. 4a). Similar observations were evident when using the STRING physical interaction network as the source of protein-protein interaction networks^30^ (t-test p = 4.8e-11 for compensation and p = 3.7e-03 for collateral loss) (Supplementary Fig. 5a).

**Figure 4.**
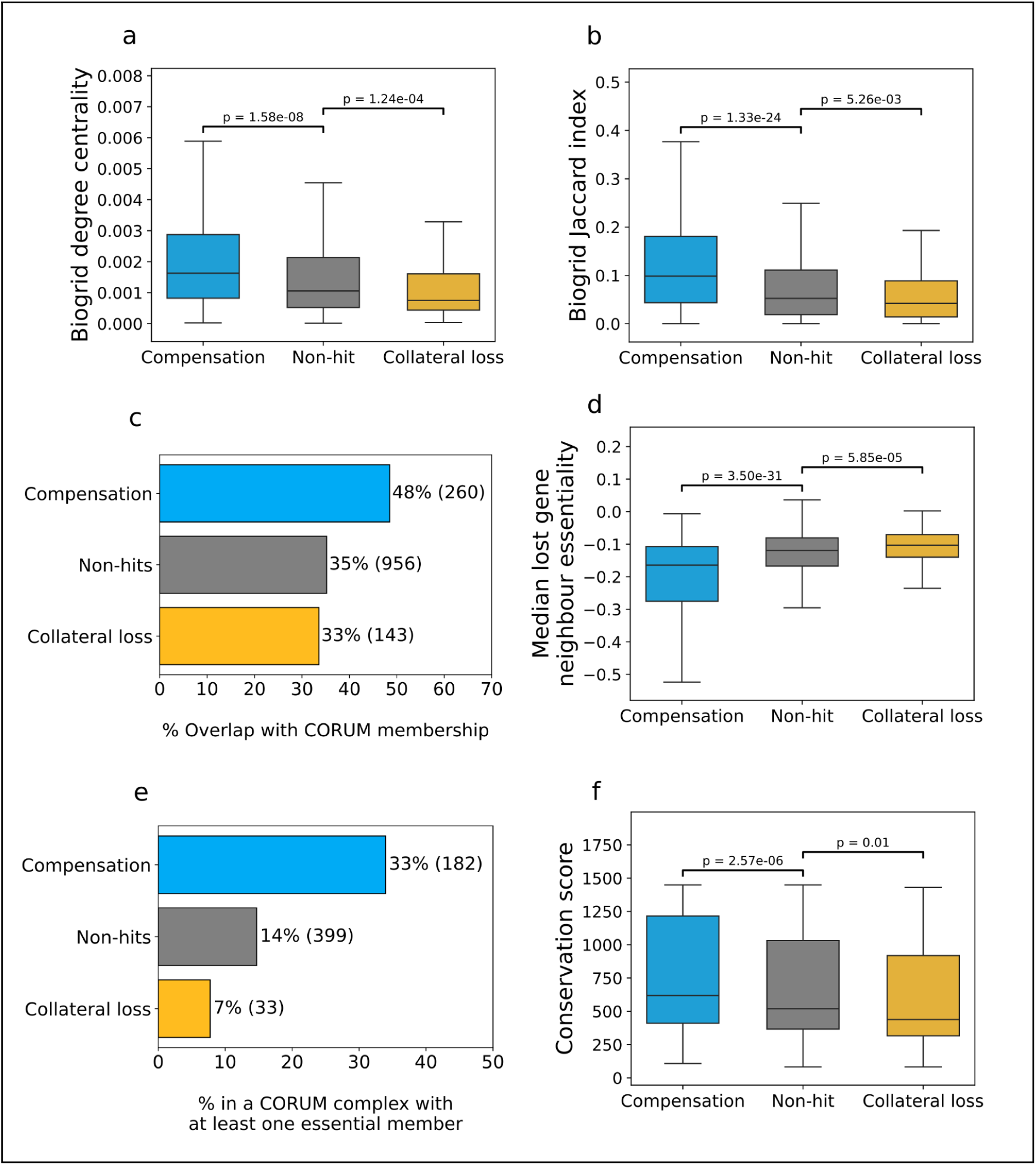
Compensation pairs are more central in the interaction network and are more conserved. **(a)** Box plots showing the degree centrality values of compensation pairs, collateral loss pairs, and non-hits, calculated using the BioGRID interaction repository ^29^. **(b)** Box plots showing the Jaccard indices of paralog pairs in all three groups, calculated using BioGRID. **(c)** Barchart showing the proportion of paralog pairs in each group where at least one paralog is a CORUM protein complex member ^31^. Counts for each group are shown in parentheses. **(d)** Box plots showing the median essentiality of the neighbours of the lost gene, for paralog pairs in all three groups, calculated using BioGRID. **(e)** Barchart showing the proportion of paralog pairs in each group where at least one paralog is a member of a CORUM protein complex with at least one broadly essential member (see Methods). Counts for each group are shown in parentheses. **(f)** Box plots showing the conservation scores of paralog pairs in all three groups, using the presence of a known ortholog in other species as a proxy for conservation across evolutionary time (see Methods). All p-values are from two-tailed t-tests (assuming equal variances).

We reasoned that proteins can only compensate for each other’s loss if they can interact with the same partners, and therefore tested whether compensatory gene pairs shared a higher proportion of their protein-protein interaction partners. The Jaccard index measures the proportion of shared interactors between two proteins when compared to the total number of interactors associated with either protein. Analysing the BioGRID database, we found that compensation hits had significantly higher Jaccard indices than non-hits (t-test p = 1.3e-24), while collateral loss hits had significantly lower Jaccard indices (p = 5.3e-03) (Fig. 4b). As before, this trend is also significant for compensation pairs when using the STRING network (p = 1.1e-07 for compensation and p = 0.65 for collateral loss) (Supplementary Fig. 5b)

Consistent with a central role in the protein-protein interaction network, we found that compensation gene pairs were significantly more likely to be annotated as belonging to protein complexes in the manually curated CORUM database^31^ (Fig. 4c). This holds both when considering either member of the pair being in a protein complex (OR = 1.7, p = 8.9e-08) or both members being in the same protein complex (OR = 2.6, p = 2.5e-06). A significant enrichment for compensation hits was also observed using an alternative source of curated protein complexes from the EBI ComplexPortal (OR = 2.1, p = 8.6e-08; Supplementary Fig. 5c) and a set of predicted protein complexes from the HuMap project (OR = 1.9, p = 1.7e-09; Supplementary Fig. 5d)^32,33^.

Compensatory relationships between paralogs may exist as a means of maintaining essential cellular functions in the face of perturbations. If that is the case, then one would expect that compensation should be enriched among genes that are involved in cellular processes whose perturbation causes a cellular growth defect. To assess this, we calculated the average growth defect (inferred from CRISPR screens in 762 cell lines, reprocessed as outlined in previous work^21^) associated with the perturbation of each protein coding gene. We then asked if the protein-protein interaction partners of genes whose loss triggered compensation generally caused a greater growth defect, and found that this is indeed the case (p = 3.5e-31) (Fig. 4d). On the other hand, genes whose loss triggered collateral loss of a paralog had significantly less essential interaction partners than non-hits (p = 5.8e-05).

Consistent with this, we find that compensation is enriched among subunits of protein complexes with at least one broadly essential subunit (CERES score < -0.6 in at least 80% of cell lines) (OR = 3.3, p = 3.5e-24)^34,35^ (Fig. 4e). In contrast, collateral loss is depleted among such complexes (OR = 0.6, p = 0.02). Genes that are broadly conserved across species tend to be involved in more essential processes. If compensation exists as a means of maintaining essential functions, then one might anticipate that compensatory gene pairs should also be more widely conserved across evolution. To test this, we assigned all paralogs a conservation score according to whether or not they had identifiable orthologs in 1,472 species (see Methods). Using this score, we found that genes involved in compensatory pairs were more widely conserved than non-hits (t-test p = 2.6e-06), while collateral loss gene pairs were less conserved than non-hits (p = 0.01) (Fig. 4f).

Overall, these results suggest that compensation pairs are more central in the protein-protein interaction network, more conserved across species, and more likely to be involved in essential functions. In contrast, collateral loss genes are more peripheral on the protein-protein interaction network, less conserved, and less likely to work in essential complexes.

### Compensation pairs are more likely to be synthetic lethal

If paralog protein compensation exists as a means of maintaining essential cellular functions in the face of perturbation, then one would anticipate that loss of one member of a compensating paralog pair would render the other paralog in the pair essential, i.e. the two genes should be synthetic lethal. This would be consistent with previous observations from budding yeast where 22 paralog pairs that exhibited compensatory relationships were found to be enriched in known synthetic lethal pairs^12^. To test this hypothesis on our set of proteomic compensation pairs we compared them to two distinct sets of synthetic lethal pairs – one set derived from genome-wide CRISPR screens in a panel of cancer cell lines from DepMap^34,35^, where loss of function of one gene could be associated with increased sensitivity to loss of its paralog^21^, and one set derived from combinatorial CRISPR screens, where paralog pairs are knocked out in combination to identify cases where the observed fitness defect of perturbing both paralogs is significantly greater than the fitness defect expected from perturbing each paralog individually^6–10,21^ (see Methods). Using either set of synthetic lethal pairs, compensation pairs were significantly enriched for synthetic lethality (DepMap: OR = 2.7; p = 7.2e-04. Combinatorial screens: OR = 2.5; p = 7.5e0-3) (Figs. 5a, 5b). Synthetic lethal hits can vary across different cell lines and screens, but we found that proteomic compensation hits were enriched in among synthetic lethal pairs whether we required them to be observed in at least one screen (OR = 2.9, p = 3.1e-08; Supplementary Fig. 6) or multiple screens (Fig. 5b).

**Figure 5.**
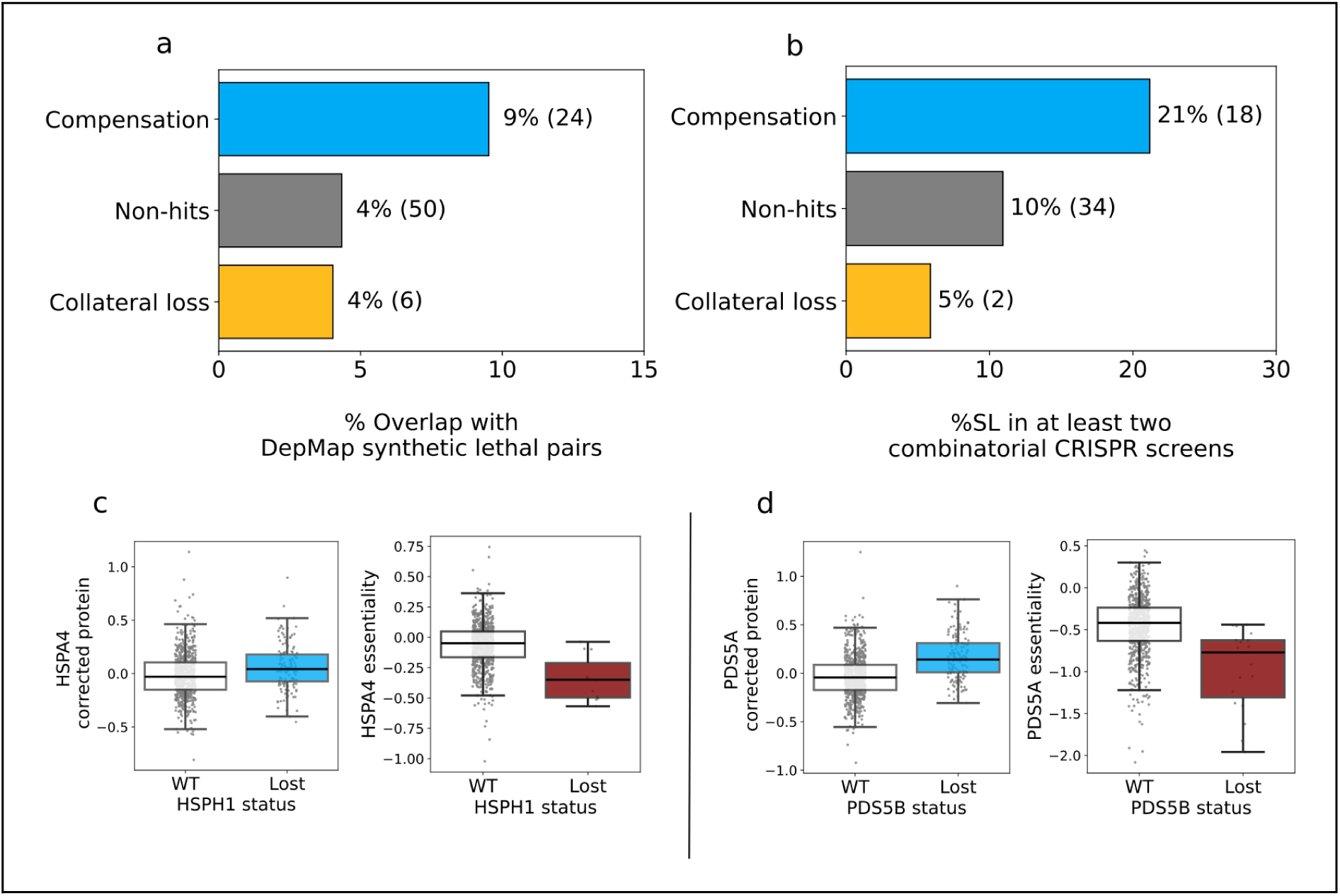
Compensation pairs are more likely to be synthetic lethal. (a, b) Barcharts showing the percentage of compensation pairs, non-hits, and collateral loss pairs which are synthetic lethal based on analysis of **(a)** DepMap data ^21^ or **(b)** based on a consensus dataset assembled from five combinatorial CRISPR screens (see Methods) ^6–10^. **(c)** Boxplots showing that HSPA4 is more abundant and more essential (i.e. lower CERES score) in cells where *HSPH1* is lost. CPTAC data was used to plot abundance and DepMap (cell line) data was used to plot essentiality. **(d)** Similar boxplots showing increase in *PDS5A* gene essentiality and PDS5A protein abundance upon *PDS5B* loss.

Synthetic lethal gene pairs that exhibit proteomic compensation include well studied pairs, such as *SMARCA2*-*SMARCA4*^36^, as well as gene pairs that have been identified as synthetic lethal in multiple screens, such as *CNOT7*-*CNOT8*, which was recently identified as the most reproducible synthetic lethality in a systematic analysis of paralog screens^37^. Among the less well characterised pairs are the paralogous heat-shock associated proteins HSPA4-HSPH1^38,39^. HSPA4-HSPH1 is a reciprocal compensation hit, i.e. *HSPH1* loss is associated with increased abundance of HSPA4 protein and *HSPA4* loss is associated with increased HSPH1 abundance. Our previous analysis of synthetic lethal relationships in a large cell line panel (DepMap) identified that cell lines that have lost *HSPH1* are more sensitive to the inhibition of *HSPA4* (Fig. 5c). An especially promising pair are the Precocious Dissociation of Sisters 5 paralogs *PDS5A* and *PDS5B,* which regulate the cohesin complex and play important roles in chromatid cohesion^40,41^. *PDS5B*, also known as *APRIN* and *AS3*, is a candidate tumour suppressor mutated or downregulated in a variety of cancer types^42–45^. Here we find that loss of *PDS5B* is associated with increased protein abundance of its paralog PDS5A (Fig. 5d). This pair was previously identified as a synthetic lethal pair in a high-throughput combinatorial CRISPR screen^8^, while our own previous analysis of single-gene knockout CRISPR screens suggested that loss of *PDS5B* was associated with increased dependency on *PDS5A* (Fig. 5d)^21^. Our results here suggest that loss of *PDS5B* is associated with an increase in the protein abundance of PDS5A and a concordant increased dependency on its encoding gene.

### Compensation and collateral loss are driven by both transcriptional and post-transcriptional regulation

We wished to assess whether compensation and collateral loss hits arose through transcriptional regulation (i.e. loss of a protein results in altered transcription of its paralog) or through post-transcriptional mechanisms (e.g. loss of a protein results in altered translation or protein degradation of its paralog). As the CPTAC dataset contains matched genomes, proteomes, and transcriptomes, we were able to run the analysis pipeline outlined in Fig. 2a on transcriptomic profiles rather than proteomic profiles in order to identify cases of transcriptional compensation and collateral loss. Furthermore, we employed a linear regression approach (similar to that previously used by Gonçalves *et al*^15^) to create a protein residuals dataset where protein abundance was adjusted for transcriptional abundance, allowing us to directly assess effects mediated by post-transcriptional regulation (see Methods; Supplementary Fig. 7). We restricted our analysis to pairs quantified in both the proteomic and transcriptomic datasets. Testing the same 4,568 paralog pairs with transcriptomic data, 607 compensation hits and 479 collateral loss hits were identified at the same FDR of 5% (Fig. 6a, Supplementary Fig. 8a). The overlap between hits observed at the transcriptomic and proteomic level was significantly more than expected by chance (OR = 12.6, p = 1.01e-131 for compensation; OR = 25.9, p = 4.4e-169 for collateral loss) but still only partial – 274 (∼48%) compensation hits identified at the protein level were not significant at the mRNA level, and 313 (∼52%) compensation hits identified at the mRNA level were not significant at the protein level (Fig. 6b). Testing the same set of pairs with the protein residual dataset, we identified 227 compensation and 140 collateral loss hits at an FDR of 5% (Fig. 6a, Supplementary Fig. 8b). As might be expected, the majority of these hits were evident at the protein level (∼90% of compensation hits and 69% of collateral loss) with a smaller overlap with the transcriptionally regulated pairs.

**Figure 6.**
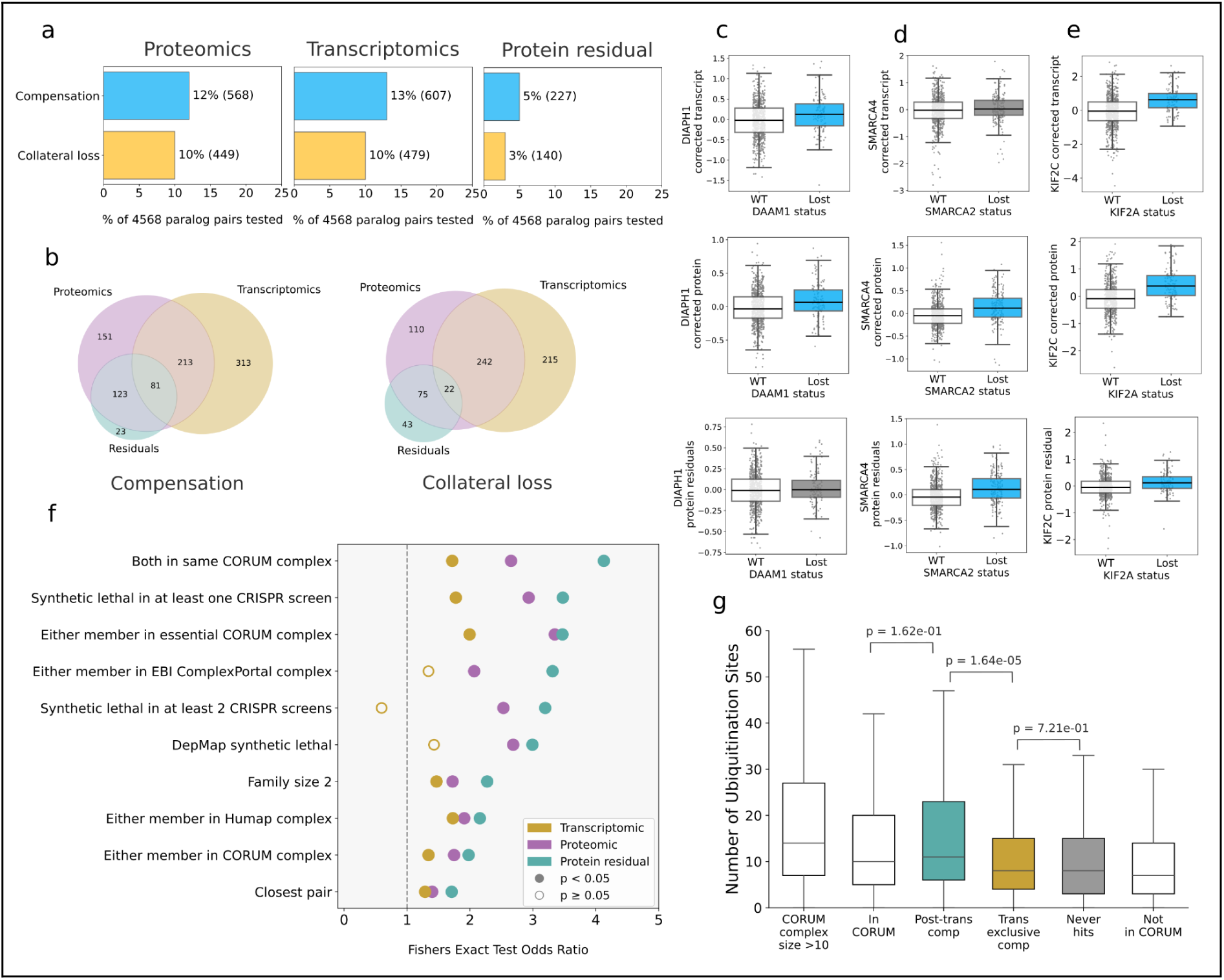
Protein compensation is regulated transcriptionally and post-transcriptionally, with post-transcriptional pairs more enriched in synthetic lethality. **(a)** Bar charts showing the number of hits identified with the proteomic dataset, transcriptomic dataset, and protein residual datasets. **(b)** Venn diagrams showing the overlap between compensation and collateral loss hits identified with the transcriptomic, proteomic, and protein residual datasets. **(c)** Boxplots showing DIAPH transcript, protein, and protein residual change associated with loss of its paralog *DAAM1*. Transcript and protein abundance shown has been corrected for lineage and self-copy number as these are covariates in the regression model. This pair is a transcriptionally-driven proteomic hit with no protein residual component. **(d)** Similar boxplots for SMARCA4-SMARCA2, a post-transcriptionally driven hit which is not a hit at the transcript level. **(e)** Similar boxplots for KIF2C-KIF2A, a pair which is a compensation hit with transcriptomic data, proteomic data, and the protein residual dataset, suggesting transcriptional and post-transcriptional regulation **(g)** Odds ratios of significant and non-significant enrichments as calculated for proteomic, transcriptomic and protein residual compensation hits. **(f)** Boxplot comparing number of ubiquitination sites between post-transcriptionally compensating paralogs (only observed in the residuals dataset), exclusively transcriptionally compensating paralogs (only observed transcriptionally), and non-compensating paralogs, with CORUM members, members of large CORUM complexes (no. of subunits >10) and non-CORUM members as controls (see Methods).

Hits identified with the transcriptomic and proteomic datasets but not the protein residual dataset likely correspond to cases of exclusively transcriptional regulation with no post-transcriptional component. For example, loss of *DAAM1* is associated with an increase in the transcript abundance and protein abundance of its paralog, DIAPH1 (Fig. 6c). DAAM1 and DIAPH1 are involved in actin polymerization and microtubule regulation^46–48^. This relationship is not detectable with the protein residual dataset, suggesting that the protein-level relationship is entirely driven by transcript-level *DIAPH1* upregulation in response to *DAAM1* loss. Hits identified with the proteomic and protein residual datasets but not the transcriptomic dataset likely correspond to cases of exclusively post-transcriptional regulation. For example, loss of the transcriptional activator *SMARCA2* is associated with an increase in protein abundance of its paralog SMARCA4, but no corresponding relationship is visible at the transcript level ^49^ (Fig. 6d). SMARCA2 and SMARCA4 are mutually exclusive members of the chromatin remodelling SWI-SNF complex ^50^. Hits identified with all three datasets likely correspond to cases of post-transcriptional regulation acting in the same direction as transcriptional regulation, resulting in an overall increase in protein abundance (a phenomenon previously observed in systematic studies of mRNA-protein correlation^51^). Compensation hits evident at both transcriptional and post-transcriptional levels include KIF2A-KIF2C, kinesin-13 proteins that regulate microtubule dynamics during mitosis^52^. Loss of *KIF2A* is associated with increased mRNA abundance of KIF2C, suggesting transcriptional regulation, and an increase in the protein residuals of KIF2C, suggesting post-transcriptional regulation also (Fig. 6e). Some pairs were identifiable only when running the analysis with the post-transcriptional dataset, corresponding to cases where an effect at the post-transcriptional level is not identifiable with proteomic data alone due to transcript-level variation. However, the majority of compensation effects detected with the protein residual dataset are evident with proteomic data.

### Post-transcriptional compensation pairs are more likely to be synthetic lethal and are more frequently ubiquitinated

We anticipated that compensatory relationships that are post-transcriptionally regulated might be more likely to involve members of protein complexes, as protein complex subunits have previously been identified as being subject to more post-transcriptional regulation ^15,16,19,53^. We therefore tested post-transcriptionally regulated pairs (i.e. protein residual compensation hits) to see if they were enriched in protein complex subunits, alongside all previously tested relationships (i.e. synthetic lethality, closest pairs, smaller paralog families). As anticipated, we found that post-transcriptionally regulated compensation pairs were more enriched than transcriptionally regulated pairs for protein complex membership, regardless of the source of complex annotations (CORUM, ComplexPortal, HuMAP) (Fig 6f). Surprisingly, we also found that post-transcriptionally regulated compensation pairs were more highly enriched in synthetic lethality than transcriptionally regulated pairs, suggesting that post-transcriptional regulation of paralog compensation is more predictive of synthetic lethality.

Post-transcriptional regulation of compensatory genes could occur via a number of mechanisms, including increased translation of the associated transcript or reduced degradation of the resulting protein products. Previous work has suggested that, for protein complex subunits in particular, altered protein degradation may be a common mechanism by which cancer cells maintain protein complex balance in the face of copy number variation of individual subunits^15,16,19^. Consistent with this, proteins with copy number variation that is attenuated at the protein level have greater numbers of ubiquitination sites^19,54^. If paralog compensation effects are also regulated by altered protein degradation, we would expect to see greater numbers of ubiquitination sites in compensatory proteins. This should be especially evident for pairs that are regulated post-transcriptionally. To test this hypothesis, we compared the numbers of ubiquitination sites for the compensating paralog in pairs only observed transcriptionally (hits only in the transcriptional dataset), pairs that were regulated post-transcriptionally (hits in the protein residual dataset), and never-hits (i.e A1s in A1-A2 pairs which were not hits in any dataset). As controls, we included proteins which are in large (>10 members) CORUM complexes (previously shown to be highly attenatued at the protein level^15,16,19^), and proteins that never feature in a CORUM complex. All sets were restricted to paralogs tested for compensation/collateral loss in our analysis, to avoid any bias in the results. In line with our expectations, proteins which are never in a CORUM complex had the lowest number of ubiquitination sites (median 7), while members of large CORUM complexes had the highest number, with a median of 14 sites (Fig. 6g). Among our hits, post-transcriptionally compensating paralogs had the highest number of ubiquitination sites (median 11), significantly higher than compensating paralogs only observed transcriptionally (median 8, Mann-Whitney U test p = 1.6e-05) (Fig. 6g). CORUM complex members overall had a median of 10 ubiquitination sites, with no significant difference from our post-transcriptionally compensating pairs (Mann-Whitney U test p = 1.6e-01). The higher number of ubiquitination sites on post-transcriptionally compensating proteins is consistent with a significant role for protein degradation in regulating paralog protein compensation.

## Discussion

Previous work has established that tumours and tumour cell lines are generally more tolerant of mutations and deletions of paralogs than singleton genes^3–5,20^. Other work has established that pairs of paralogs are frequently synthetic lethal, suggesting that they can directly compensate for each other’s loss^6–10^. How this compensation works at a molecular level has not been systematically explored in the context of cancer or indeed in any human cells. Here, we have used proteomic profiles of isogenic cell lines and tumour samples to understand how loss of one gene impacts the protein abundance of its paralog. We found evidence of relatively frequent active compensation between paralogs as well as collateral loss, with the former being more common. Our subsequent analyses of the identified pairs suggest that compensatory relationships are more likely to be observed among pairs that are well connected on the protein-protein interaction network, highly conserved across species, and involved in essential functions. In contrast, collateral loss is observed for pairs that are less well conserved, more peripheral on the protein-protein interaction network, and less connected to essential processes. Compensatory relationships, unlike collateral loss relationships, are more likely to occur between pairs that are known to exhibit synthetic lethality. Taken together, our results suggest a model whereby compensatory relationships stabilise essential protein-protein interaction subnetworks or complexes in the face of genetic perturbation in cancer. Following loss of one gene, the protein abundance of a highly sequence similar paralog is increased, and this protein takes the place of the lost gene in the protein-protein interaction network. Cancer cells then become dependent on this protein for survival (Fig. 7). Previous work has identified individual examples consistent with this model, for example loss of the cohesin subunit STAG2 results in increased protein abundance of its paralog STAG1 and an increased dependency upon *STAG1*; our results suggest that it this may be a relatively common phenomenon^55,56^.

**Figure 7.**
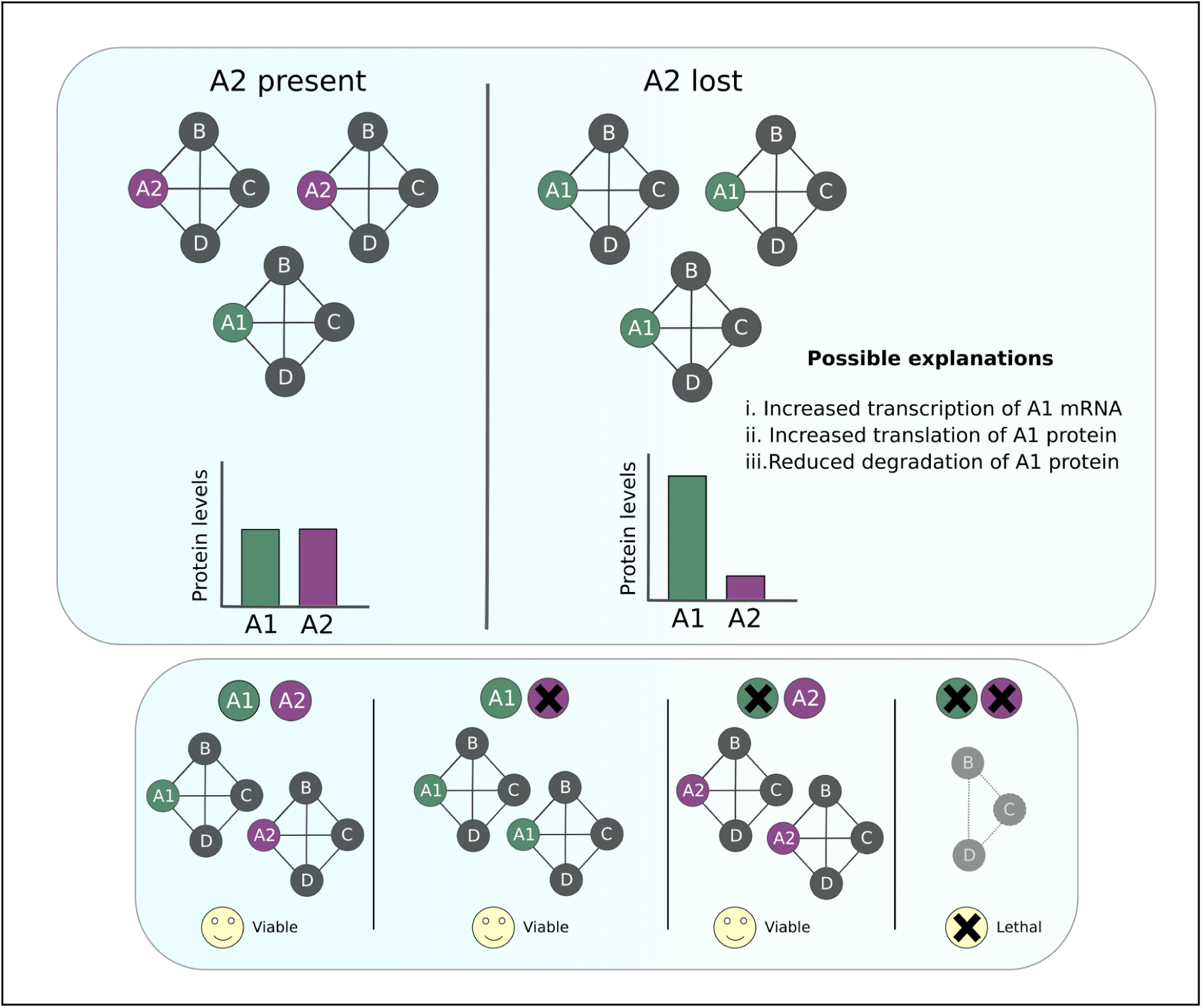
Proposed model for protein-protein interaction network rewiring leading to proteomic compensation and synthetic lethality. Our results are consistent with a model where, for pairs with post-transcriptional compensation effects, increased incorporation of A1 into A2’s protein interactions following A2 loss results in an increase of its abundance through reduced degradation of A1 subunits. A1 becomes essential in backgrounds where A2 is lost, i.e. the pairs are synthetic lethal.

### Mechanisms of compensation and collateral loss

Compensatory relationships could occur through a number of mechanisms, including transcriptional regulation or post-transcriptional regulation. Our analysis suggests that both types of regulation contribute to compensation effects observed at the level of total protein abundance.

Considering only regulation of mRNA abundance, a diversity of mechanisms could be employed. Perhaps most obviously, paralogous transcription factors could directly regulate each other’s transcription. However, alternative means of regulating mRNA abundance also exist. For example, in mice, the ribosomal paralog *RPL22* regulates the protein abundance of its paralog *RPL22L1* by directly binding and destabilising *RPL22L1* mRNA ^57^. Also in mice, MBNL1, an RNA binding protein, regulates the splicing of its paralog *MBNL2* such that loss of *MBNL1* results in the altered inclusion of specific exons and a more stable protein^58^. An alternative mechanism, termed genetic compensation or transcriptional adaptation, has recently been identified whereby nonsense mutation of one paralog may trigger an increase in the transcription of its paralog. It is not fully clear how this increased transcription occurs, but the mechanism appears to be dependent on the nonsense-mediated decay pathway and so is unlikely to be relevant for pairs triggered solely by copy loss^59,60^.

Previous work has established that post-transcriptional regulation is an important mechanism by which cancer cells adapt to genetic perturbation and has primarily focused on protein degradation. In particular, multiple studies have found that copy number variation in cancer often results in concordant effects on mRNA abundance but that this is frequently attenuated at the protein level, i.e. a doubling of copy number may lead to a similar increase in mRNA abundance but a much smaller increase in protein abundance^15–17^. Genes subject to this attenuation are significantly more likely to encode subunits of protein complexes and also have more known ubiquitination sites, suggesting potential regulation via protein degradation^15,16,19^. Other work has established that many protein complex subunits are produced in excess, with those proteins not incorporated into complexes subsequently degraded, providing a means of maintaining balance between subunits^61–64^. Our finding that paralog compensation pairs, especially those that are post-transcriptionally regulated, are enriched in protein complex subunits and more highly ubiquitinated suggests a potentially similar model. If both members of a paralog pair compete for membership of the same protein complex, as is the case for many pairs, then any orphan subunits not incorporated into a complex will be degraded. When one paralog has reduced abundance, resulting from mutation or deletion, its counterpart will have increased incorporation into the complex, reduced degradation, and so increased abundance. This form of compensation could occur without any need for altered transcription. Further experiments will be required to disentangle the relative contributions of different regulatory mechanisms for compensatory pairs.

We do not have a simple model for collateral loss pairs. The simplest model, supported by systematic work in yeast, would be that pairs of paralogs often form heterodimers and hence stabilise each other. When one paralog is lost, the other loses stability and is then degraded. The strongest effect we see, GMPPA/GMPPB, corresponds to a pair known to form a heterodimer, consistent with this model. Since there is considerable overlap between collateral loss hits identified at the transcript level and those identified at the protein level, it is also likely that many of these relationships involve some transcriptional co-regulation. However, this is unlikely to explain all collateral loss pairs and so additional mechanisms need to be identified.

### Proteomic compensation as a predictor of synthetic lethality

We found that pairs that exhibit proteomic compensation are enriched among gene pairs with a known synthetic lethal relationship. Previous work has identified a number of synthetic lethal relationships between pairs that exhibit proteomic compensation, including *STAG1*/*STAG2* and *SMARCA2*/*SMARCA4*. Our finding that there is a general relationship between the two is consistent with earlier reports in yeast, where 22 pairs that displayed compensatory relationships were found to be enriched in known synthetic lethal pairs^12^. As the majority of paralog pairs have yet to be experimentally tested for synthetic lethality, the hundreds of protein compensation pairs we have identified represent candidates worth prioritising for future study. As these involve genes recurrently lost in cancer, they may be especially promising from a therapeutic point of view. Among the gene pairs with a compensatory relationship, and a known synthetic lethality, *PDS5A*/*PDS5B* appears especially promising as a therapeutic target— there is evidence of synthetic lethality both in a large scale analysis of diverse cell lines^21^ and in a combinatorial screen^8^, and *PDS5B* is recurrently deleted in patient tumours.

### Why do active protein compensation relationships exist?

The compensatory relationships we observe here are only evident in the face of a genetic perturbation – loss or mutation of one paralog triggers an increase in the abundance of another. It is extremely unlikely that these compensatory relationships have evolved as a means of protecting cells against future, unseen genetic perturbations such as those that occur in cancer. One potential explanation, proposed for compensatory relationships observed in yeast, is that these regulatory relationships exist as a means of buffering cells against variation in gene expression so that the total abundance of a paralog pair is maintained despite fluctuations in the abundance of each individual paralog ^65^. In this way the compensatory relationships we observe in cancer, that contribute to genetic robustness, may simply reflect existing regulatory relationships that have evolved to protect cells against stochastic and environmental variation.

## Methods

### Strategy for selection of paralog pairs for HAP1 knockouts

17 paralog pairs were chosen to be knocked out in a HAP1 background to measure the effect of the loss of a paralog on the other member of a pair. From a candidate set of genes that are non-essential in the HAP1 background and detectable in the HAP1 transcriptome, we selected pairs that had a known or predicted synthetic lethal relationship (Tables 1 and 2).

**Table 1.** Rationale for choosing pairs to knockout in HAP1 background. Synthetic lethality is defined as being synthetic lethal either in any background in at least one out of five combinatorial CRISPR screens, or being a gene pair where the loss of one gene results in an increase in essentiality of the other gene (based on DepMap data) ^6–10,21^. Protein complex membership is defined as either member of the pair being annotated as a member of a CORUM complex ^31^.

| Gene pair | Synthetic lethal | Synthetic lethal in | Protein complex members |
| --- | --- | --- | --- |
| *CHMP1A-CHMP1B* | Yes | Thompson et al,<br>Parrish et al <sup>8,10</sup> | Yes |
| *SMARCC1-*<br>*SMARCC2* | No | NA | Yes |
| *DDX17-DDX5* | Yes | Ito et al <sup>9</sup> | Yes |
| *CDK19-CDK8* | No | NA | Yes |
| *MBD2-MBD3* | Yes | De Kegel et al <sup>21</sup> | Yes |
| *CREBBP-EP300* | Yes | De Kegel et al,<br>Parrish et al,<br>Gonatopoulos-Pourn<br>atzis et al, Ito et al<br><sup>7-9,21</sup> | Yes |
| *SEC31A-SEC31B* | No | NA | No |
| *SEC23A-SEC23B* | Yes | Thompson et al,<br>Parrish et al,<br>Gonatopoulos-Pourn atzis et al<sup>7,8,10</sup> | No |
| *RAB1A-RAB1B* | Yes | Gonatopoulos-Pourn<br>atzis et al <sup>7</sup> | No |
| *VPS4A-VPS4B* | Yes | De Kegel et al, Ito et<br>al <sup>9,21</sup> | No |
| *ARFGAP2-*<br>*ARFGAP3* | No | NA | No |
| *KAT2A-KAT2B* | Yes | De Kegel et al,<br>Thompson et al, Ito<br>et al <sup>9,21</sup> | Yes |
| *ARID1A-ARID1B* | Yes | De Kegel et al,<br>Thompson et al,<br>Parrish et al,<br>Gonatopoulos-Pourn<br>atzis et al, Ito et al<br><sup>7-10,21</sup> | Yes |
| *SMARCA2-*<br>*SMARCA4* | Yes | De Kegel et al,<br>Thompson et al, Ito<br>et al <sup>9,10,21</sup> | Yes |
| *MAGT1-TUSC3* | Yes | Gonatopoulos-Pourn<br>atzis et al <sup>7</sup> | Yes |
| *COPS7A-COPS7B* | Yes | Dede et al <sup>6</sup> | Yes |
| *ASF1A-ASF1B* | Yes | Parrish et al <sup>8</sup> | Yes |

**Table 2.** Catalogue IDs for all proteomically profiled cell lines. All cell lines were obtained from Horizon Discovery.

| **Cell line** | **Catalogue number** |
| --- | --- |
| HAP1_WT_KO | C631 |
| HAP1_ARFGAP2_KO | HZGHC84364 |
| HAP1_ARFGAP3_KO | HZGHC26286 |
| HAP1_ARID1A_c003_KO | HZGHC000618c003 |
| HAP1_ARID1A_c010_KO | HZGHC000618c010 |
| HAP1_ARID1B_KO | HZGHC000582c007 |
| HAP1_ASF1A_KO_clone4 | HZGHC007215c004 |
| HAP1_ASF1A_KO_clone5 | HZGHC007215c005 |
| HAP1_ASF1B_KO | HZGHC55723 |
| HAP1_CDK8_KO | HZGHC000325c001 |
| HAP1_CDK19_KO | HZGHC000078c020 |
| HAP1_COPS7A_KO | HZGHC000914c001 |
| HAP1_COPS7B_KO | HZGHC000636c011 |
| HAP1_CHMP1A_KO | HZGHC005340c003 |
| HAP1_CHMP1B_KO | HZGHC007219c010 |
| HAP1_CREBBP_KO | HZGHC001109c005 |
| HAP1_EP300_KO | HZGHC001318c011 |
| HAP1_DDX5_KO | HZGHC006136c012 |
| HAP1_DDX17_KO | HZGHC007221c009 |
| HAP1_KAT2A_KO | HZGHC000929c001 |
| HAP1_KAT2B_KO | HZGHC000948c012 |
| HAP1_KMT2C_KO | HZGHC003909c012 |
| HAP1_MAGT1_KO | HZGHC84061 |
| HAP1_TUSC3_KO | HZGHC004076c007 |
| HAP1_MBD2_KO | HZGHC003628c010 |
| HAP1_MBD3_KO | HZGHC001113c009 |
| HAP1_RAB1A_KO | HZGHC007227c002 |
| HAP1_RAB1B_KO | HZGHC001225c011 |
| HAP1_SEC23A_KO | HZGHC001218c001 |
| HAP1_SEC23B_KO | HZGHC020241c010 |
| HAP1_SEC31A_KO | HZGHC22872 |
| HAP1_SEC31B_KO | HZGHC25956 |
| HAP1_SMARCA2_KO | HZGHC004055c012 |
| HAP1_SMARCA4_KO | HZGHC002878c004 |
| HAP1_SMARCC1_KO | HZGHC002476c011 |
| HAP1_SMARCC2_KO | HZGHC002783c011 |
| HAP1_VPS4A_KO | HZGHC004623c005 |
| HAP1_VPS4B_KO | HZGHC006889c011 |

### Proteomic profiling of HAP1 knock-out (KO) cell lines

For protein abundance experiments, samples were collected in lysis buffer containing 6 M Guanidine-Hydrochloride (Gu-HCl), 200mM Hepes pH 8.5, 1mg/ml Chloracetamide,0 and 50mM TCEP (Tris(2-carboxyethyl)phosphine hydrochloride. Cell pellets were probe sonicated and boiled at 95°C for 5 min. Cell lysates were clarified by centrifugation (20 000 g, 5 min). HAP1 KO cell lines were lysed and digested in 3 batches. Batch 1 consists of WT, ARFGAP2_KO, ARFGAP3_KO, ARID1A_c003_KO, ARID1A_c010_KO, ARID1B_KO, ASF1A_C4_KO, ASF1A_C5_KO, ASF1B_KO, CDK8_KO, CDK19_KO, CHMP1A_KO, and CHMP1B_KO cell lines. Batch 2 consists of WT, COPS7A_KO, COPS7B_KO, DDX5_KO, DDX17_KO, CREBBP_KO, EP300_KO, KAT2A_KO, KAT2B_KO, KMT2C_KO, MAGT1_KO, and TUSC3_KO cell lines. Batch 3 consists of WT, MBD2_KO, MBD3_KO, RAB1A_KO, RAB1B_KO, SEC23A_KO, SEC23B_KO, SEC31A_KO, SEC31B_KO, SMARCA2_KO, SMARCA4_KO, SMARCC1_KO, SMARCC2_KO, VPS4A_KO, and VPS4B_KO cell lines.200 µg of proteins were digested with LysC (New England Biolabs) by adding 0.4 µg LysC (Promega) per sample for 4 hr at 37°C. Samples were diluted to 1 M Gu-HCl and then digested with Trypsin for 16 hr at 37°C. Trypsin activity was inhibited by acidification of samples to a concentration of 1% trifluoroacetic acid. Digests were clarified by centrifugation (20,000 g, 5 min), samples were desalted on a C18 Stage tip, and eluates were analysed by HPLC coupled to a Q-Exactive Plus mass spectrometer as described previously but with an extended gradient of 120 min ^66^. Peptides and proteins were identified and quantified with the MaxQuant software package (2.6.4.0), and label-free quantification was performed by MaxLFQ ^67^. The search included variable modifications for the oxidation of methionine, protein N-terminal acetylation, and carbamidomethylation as fixed modification. The false discovery rate, determined by searching a reverse database, was set at 0.01 for both peptides and proteins. Proteins identified were filtered to remove reverse sequences and potential contaminants. LFQ intensity values were log2-transformed to normalize the data distribution. A mean centering procedure was applied to adjust for systematic differences between samples. This involved calculating sample-specific adjustment factors based on the difference between each sample’s mean and the global mean. These factors were then subtracted from all LFQ intensities Proteins quantified in fewer than 20% of the samples were dropped. Missing values were imputed in each sample with the lowest protein measurement for that sample.

### Differential expression of paralogous proteins upon knockout of the other paralog

A two-tailed equal variance t-test was performed for every HAP1 clone to compare the abundance of the knocked-out protein in the knockout cell lines vs. its abundance in the wild-type cell lines. Four biological replicates were profiled for every knockout while the wild-type had twelve biological replicates. Out of 34 candidate genes, this test could not be performed for 6 knockouts as their cognate proteins were not measured in any of the HAP1 wild-type samples. Tests were also not performed for ASF1A/ASF1B since MaxQuant failed to calculate unique abundance values for each protein (due to all measured peptides being shared between both paralogs). Following multiple testing correction using the Benjamini-Hochberg method, clones without significantly (FDR < 5% and uncorrected p-value < 0.05) lower protein abundance in the knockout were dropped from the analysis (3 out of 26 clones). For the 23 clones with validated knockouts, a similar t-test was performed to compare the abundance of each knocked out gene’s paralogs. This was done for all paralogs that were measured in at least one sample. In addition to assessing paralog pairs with symmetric knockouts, i.e. A1-A2 pairs where we had knockouts for both A1 and A2, we also compared the levels of other paralogs in the family (see below for paralog selection criteria), e.g. In the case of the pair RAB1A-RAB1B, we compared RAB1A abundance in *RAB1B* knockout (KO) vs wild-type (WT) and RAB1B abundance in *RAB1A* KO vs WT. We also compared the abundances of RAB18 and RAB35 in both these knockouts with the wild types. In this way, a total 46 t-tests were performed (Fig. 1). All t-tests were performed with 4 samples in the knockout group and 12 samples in the wild type group, i.e. 14 degrees of freedom.

### Detecting paralog compensation and collateral loss in primary tumours

The CPTAC PanCan datasets, containing collated and normalized proteomics and transcriptomics data for samples from 9 CPTAC studies, were used for this analysis ^71^. In total, the datasets had 1023 samples, with each study representing a different tissue type. Copy number data was obtained from the PanCan CPTAC GISTIC dataset, which contained absolute gene-level copy number values. Ovarian cancer samples were dropped since this study had fewer than 100 samples.

### Selection criteria for paralog pairs

A symmetric set of 83,404 paralog pairs was obtained from Ensembl release 93 ^72^. Pairs with sequence identity <= 0.2 and family size >= 20 were excluded. Paralog pairs were excluded when either member of the pair was measured in fewer than half the samples present in the dataset. To minimize the possible confounding effect of co-regulation or linkage, paralog pairs were excluded if both members of the pair were located on the same chromosome (Supplementary Fig. 2).

### Calling gene loss

Gene losses were annotated for every sample within each dataset. Absolute copy number values called using the GISTIC algorithm were used to annotate hemizygous deletions (i.e. copy number of -1 in a given sample instead of a copy number of 0) ^73^. Gene losses were annotated for all genes that were members of suitable paralog pairs (as determined in the previous section). Genes were then filtered to a set that was hemizygusly lost in at least 20 samples. Genes were dropped when their protein abundance was not quantified in at least 20 of the samples where they were hemizygously lost. Gene losses were only considered to be valid if they were associated with a significant drop in the abundance of their cognate protein. To calculate this, a linear model was fit using ordinary least squares with protein abundance as the dependent variable, gene loss as the independent variable, and tissue-of-origin/study (as each CPTAC study corresponded to a different cancer type) for the sample as a covariate. The two-tailed p values for the t-statistic of the gene loss variable in each model was calculated and nominally significant proteins with a p-value below 0.05 were retained. In this way, we identified cases where the loss of a gene affected its protein abundance. Paralog pairs where at least one member was lost in a minimum of 20 samples, with this loss being associated with a significant drop in protein abundance, were retained.

### Generation of protein residual datasets

A ‘protein residual’ dataset was generated to identify relationships between paralogs that arose solely post-transcriptionally. This was done by regressing out self-transcript abundance from protein abundance for each sample. In order to appropriately account for the effect of tissue-of-origin, which may impact mRNA and protein abundance separately, linear models were built for each gene to explain mRNA abundance and protein abundance (separately) with the study/tissue type variable. The residuals of these models, i.e. study corrected mRNA and study-corrected protein, were used to obtain protein residuals. This was done by fitting linear models to explain study-corrected protein abundance using study-corrected mRNA abundance and taking the residuals. (Supplementary Fig. 7).

### Paralog compensation and collateral loss analysis

Gene losses were annotated in every sample and paralog pairs were filtered to a testable set as detailed above. For every paralog pair A1-A2, the dataset was partitioned into a set where A2 had been hemizygously lost, and a set where two copies of A2 were present (excluding samples with amplifications or homozygous deletions). A linear model was then fit to explain protein abundance of the other paralog, A1, based on A2 loss (as a binary variable), cell line lineage/tumour type, and A1 copy number. Lineage was included as a covariate to minimize the confounding effect of tissue-specific expression patterns. A1’s own copy number was also included as a covariate to avoid misattributing variation caused by self-copy number changes to A2 loss. In total, 4,568 pairs were tested when calling loss based on hemizygous copy number loss. Two-tailed p values for the t-statistic of the A2 loss variable in each model was extracted, and following Benjamini-Hochberg correction for multiple testing, any pairs with False Discovery Rate (FDR) for A2 loss < 5% and uncorrected p-value < 0.05 were considered to be hits-either compensation if A2 loss was associated with increased A1 abundance, or collateral loss if A2 loss was associated with decreased A1 abundance. The analysis was run separately for the proteomic, transcriptomic, and protein residual datasets detailed above. Lineage and self-copy number were not included as covariates for the protein residual regressions as these variables had already been regressed from the data.

### Assessing the enrichment of collateral loss / compensation using binary variables

Several factors were assessed for enrichment in the sets of compensation and collateral loss hits (separately, as compared to non-hits). Only unique pairs were retained for this analysis. These included synthetic lethality, protein complex membership of either paralog, membership of both paralogs to the same protein complex, whether or not the pair is a ‘closest pair’, i.e. the members of the pair are each other’s most sequence-similar paralogs and whether or not any other paralogs belong to the family. Synthetic lethal gene pairs from combinatorial screens were obtained from a consensus dataset previously generated by De Kegel et al, containing the results of four combinatorial CRISPR screens ^6–8,10,21^. This was augmented with hits from another combinatorial CRISPR screen ^9^. To ensure only synthetic lethal pairs, rather than negative genetic interaction pairs, were retained from Ito et al we filtered based on log fold change as performed for the Gonatopoulos-Pournatzis dataset in De Kegel et al. An LFC threshold of -0.8 was used for the Ito et al screen. A separate set of synthetic lethal gene pairs generated by De Kegel et al by identifying increases in essentiality of a gene upon loss of its paralog was also used ^21^. Protein complex membership was annotated based on data from CORUM, EBI ComplexPortal, and HuMap (complexes with confidence between 1-3) ^31–33^. Sequence identity, and family size annotations were obtained from Ensembl 93 ^72^. Pairs were also annotated based on whether they were compensation or collateral loss hits in the other direction, if they were testable in this direction. For the latter tests, pairs were not filtered to unique pairs, i.e. A1-A2 was a distinct observation from A2-A1. For both sets of hits (i.e. compensation and collateral loss), a series of Fishers Exact Tests were performed to check whether a given factor was significantly (FDR < 5% following Benjamini-Hochberg multiple testing correction of the Fishers Exact Test p-values) enriched in the set of hits. All odds ratios reported are unconditional maximum likelihood estimates calculated using the Scipy stats function fishers_exact. Supplementary Table 9 includes the total number of overlapping pairs included for each test.

### Assessing the enrichment of collateral loss / compensation pairs using quantitative variables

T-tests (two-tailed, assuming equal variance) were used to assess whether compensation pairs and collateral loss pairs had significantly different sequence identities, family sizes, neighbour essentialities (i.e. mean fitness defect associated with deletion of the interaction partners of the lost gene), protein-protein interaction network degree centralities, and Jaccard Indices compared to non-hit pairs (FDR < 5% following Benjamini Hochberg multiple testing correction). Sequence identity and family size were obtained from Ensembl 93. Essentiality data was obtained from DepMap 20Q4 (CERES scores) reprocessed as outlined in previous work ^21^. Jaccard indices were calculated separately from a network representation of the Biogrid database filtered to exclude genetic interaction data and from the STRING physical subnetwork (unweighted edges). The STRING and Biogrid databases were also used to compute degree centrality, shared interactors, and total interactors. Phylogenetic conservation scores correspond to the number of species possessing at least one known ortholog among 1472 eukaryote species in the orthology database OrthoInspector (Eukaryota2023). Ortholog relationships for human proteins were retrieved using the OrthoInspector REST API with their Uniprot accession identifiers ^74^. Ensembl version 111 was used to calculate these scores ^75^. T-statistics and degrees of freedom (comp_vs_non_df, cl_vs_non_df) for all t-tests run are recorded in Supplementary Table 10.

### Ubiquitination sites

Data on ubiquitination sites for each protein was obtained from PhosphoSitePlus v 6.7.4 ^76^. Proteins not present in this dataset were annotated as having 0 ubiquitination sites. Two-tailed Mann-Whitney U tests were performed to assess differences between groups (Groups shown in Fig. 6g). Sample size was 502 for the post-transcriptional compensation - transcriptomic-exclusive compensation test, and 651 for the CORUM member proteins - post-transcriptional compensation test.

### Computational tools used

All data analysis and visualization was performed using Python (v3.8.10), utilizing the following libraries: NumPy (v1.24.4), Pandas (v2.0.3), Pandarallel (v1.6.5), TQDM (v4.66.4), Statsmodels (v0.14.1), SciPy (v1.10.1), NetworkX (v3.1), Matplotlib (v3.7.4), Seaborn (v0.13.2), and matplotlib-venn (v0.13). Figures were created using Inkscape v1.3.

## Data availability

Data generated in this study, i.e. the HAP1 mass spectrometry proteomics data, have been deposited to the **ProteomeXchange Consortium** via the PRIDE partner repository with the dataset identifier **PXD055643** and **10.6019/PXD055643** ^68^. CPTAC PanCan data analyzed in this study is accessible from the Proteomic Data Commons (https://proteomic.datacommons.cancer.gov/pdc/cptac-pancancer) (Proteomics: Proteome_BCM_GENCODE_v34_harmonized_v1.zip-all tumour datasets concatenated and Ensembl IDs mapped to gene symbols using HUGO Gene Nomenclature Committee, i.e. HGNC mapping ^69^. Gene-level RNAseq files from tumour datasets and gene-level GISTIC copy number calls for tumor datasets) ^70^.

## Code availability

All code used to run this analysis and to generate all figures and supplementary tables is available at https://github.com/cancergenetics/proteomic_paralog_compensation.

## Acknowledgements

This work was supported by Science Foundation Ireland (under grant number 20/FFP-P/8641 and 18/CRT/6214) and the Irish Research Council (2017/2018 Laureate Award).

## Author Contributions

Conceptualization: A.V. and C.J.R.; Formal analysis: A.V., S.R.U., B.D.K., O.D.; Data curation: A.V.; Funding Aquisition: C.J.R.; Investigation: N.Q., A.B.C., K.W., A.V.K., T.L. Supervision: C.J.R.; Writing – original draft: A.V., C.J.R.; Writing – review and editing: B.D.K., T.L.

