## Supplementary Figures for "Paralog protein compensation preserves protein-protein interaction networks following gene loss in cancer"

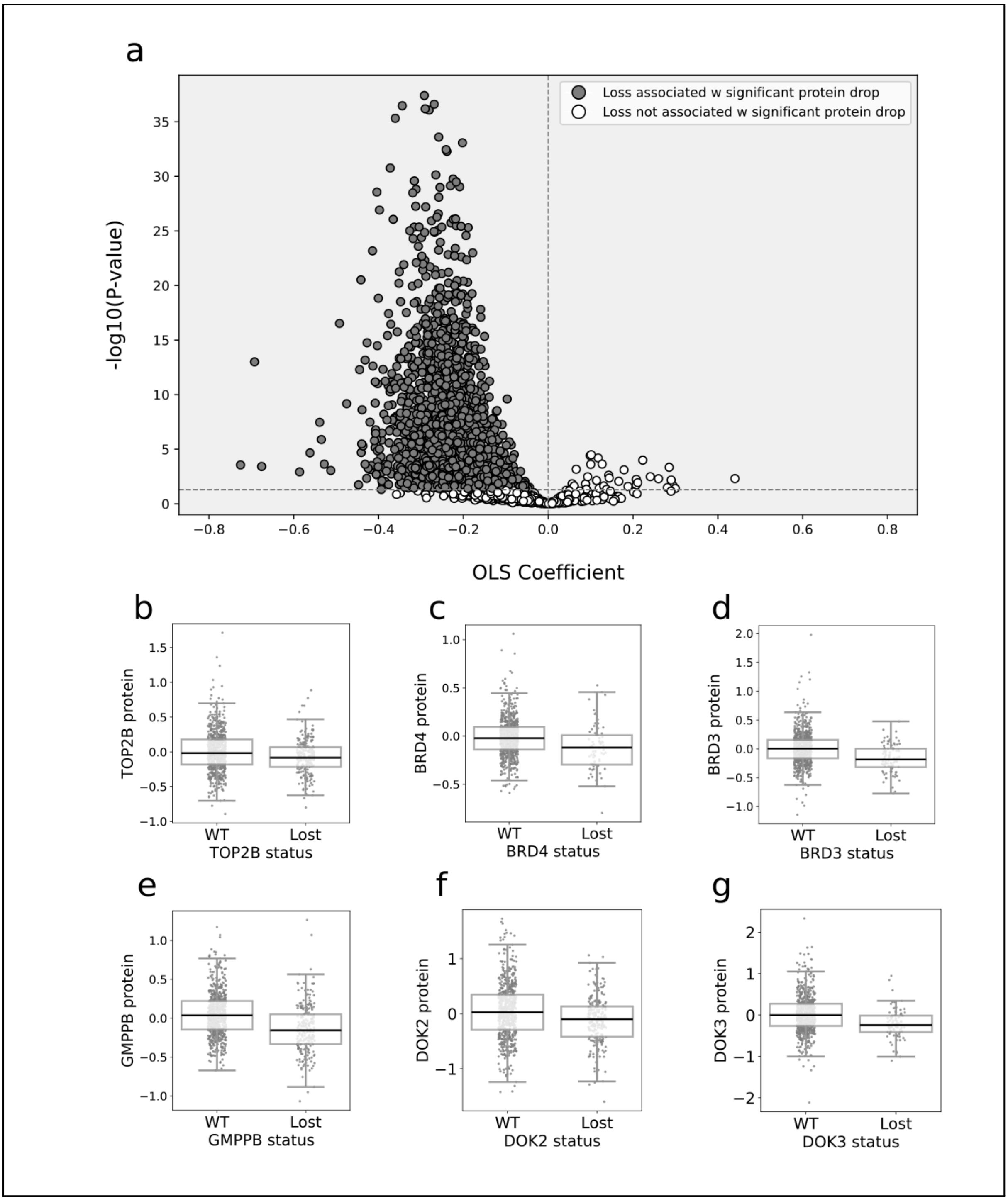


**Supplementary Figure 1. Most hemizygous losses are associated with a drop in protein abundance. (a)** Volcano plot showing decrease in protein abundance of most tested proteins following hemizygous loss (using CPTAC data) (**b-g)** Boxplots comparing abundance of the 'lost' paralog in samples where it is hemizygously lost versus samples retaining both copies of the gene, for all pairs shown in Figure 2 (*TOP2B*, *BRD4*, *BRD3*, *GMPPB*, *DOK2*, *DOK3*).

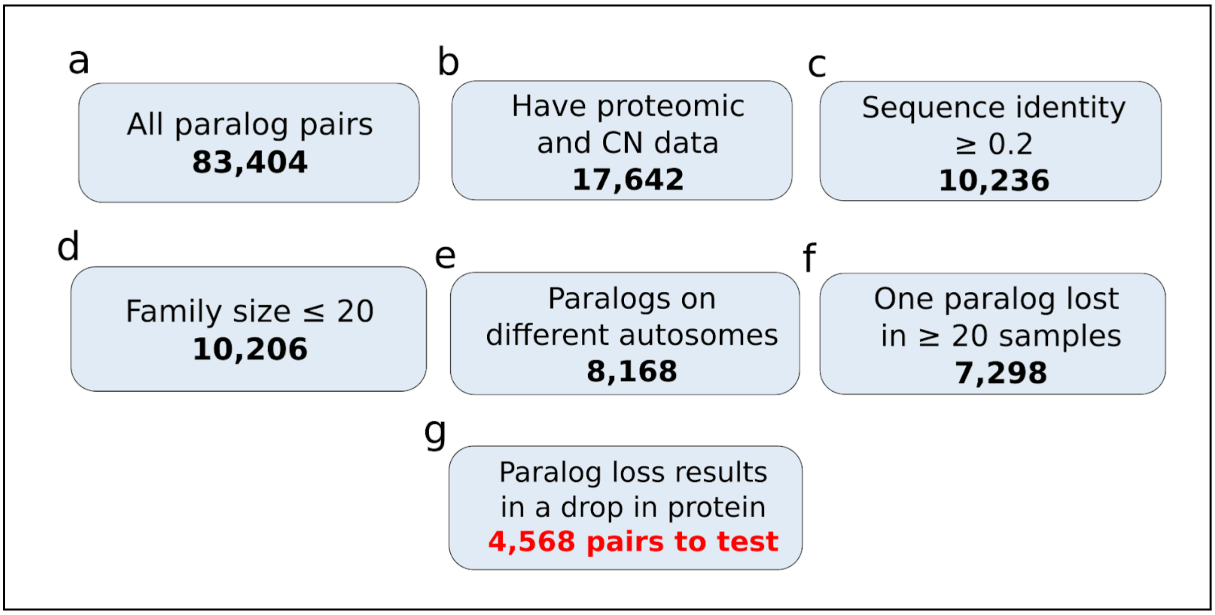


**Supplementary Figure 2. Workflow diagram showing the number of paralog pairs filtered out at each step.** 4,568 paralog pairs are tested in the final analysis of CPTAC proteomic data.

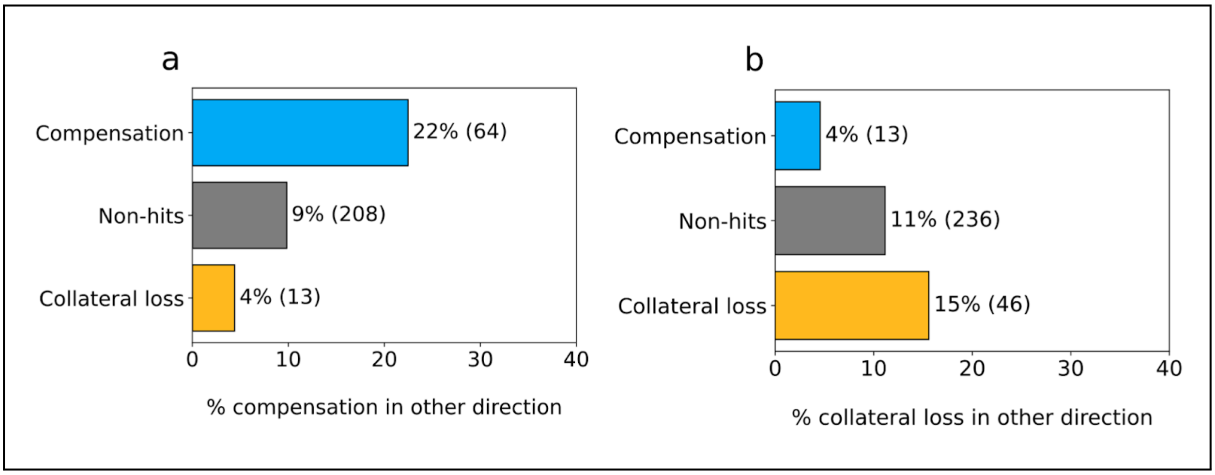


**Supplementary Figure 3. Paralog pairs tested in both directions are enriched for reciprocal hits (a)** Barchart showing the percentage of pairs in the compensation, non-hit, and collateral loss groups that are compensation hits in the other direction. **(b)** Similar barchart showing the percentage of pairs in each set that are collateral loss hits in the other direction. Only pairs tested in both directions are considered here.

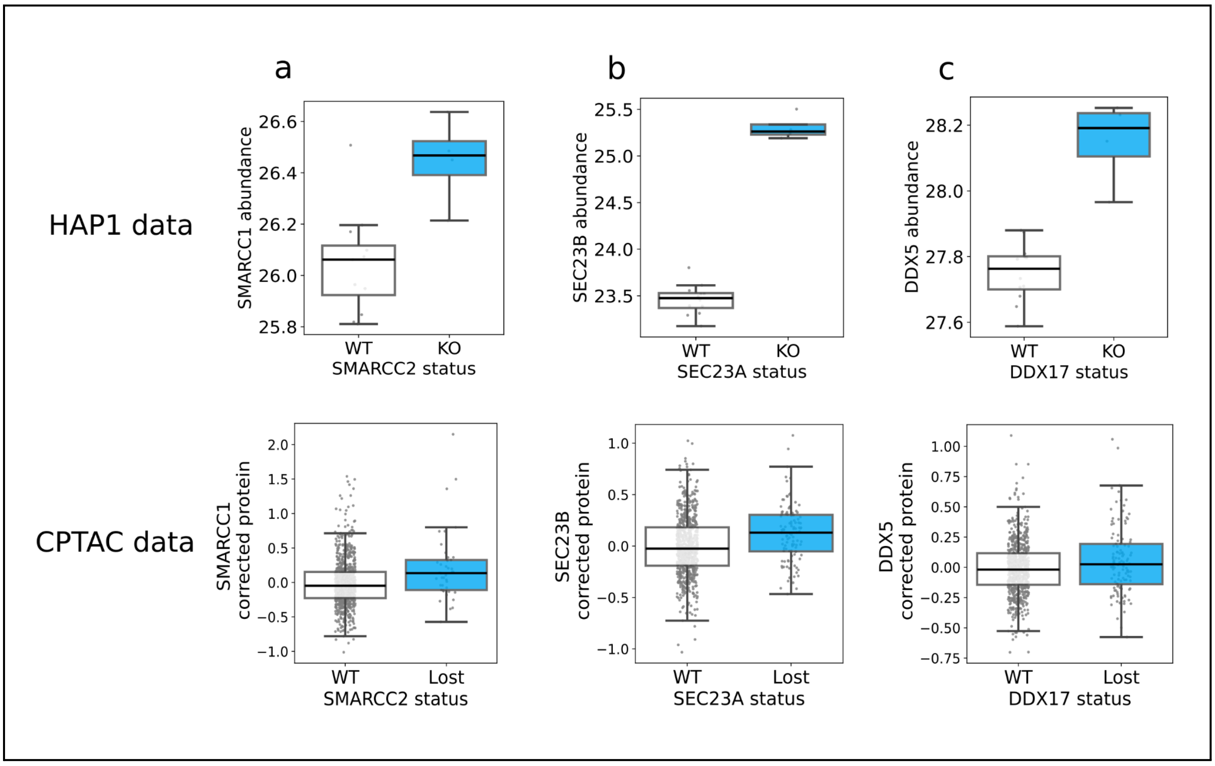


**Supplementary Figure 4. Three paralog pairs are compensation hits both with the HAP1 cell line dataset and the CPTAC tumour sample dataset. (a)** Increased SMARCC1 protein abundance is associated with *SMARCC2* knockout and *SMARCC2* loss in tumour samples. **(b)** Similar plots showing SEC23B abundance increase associated with *SEC23A* loss. **(c)** Similar plots showing DDX5 abundance increase associated with *DDX17* loss.

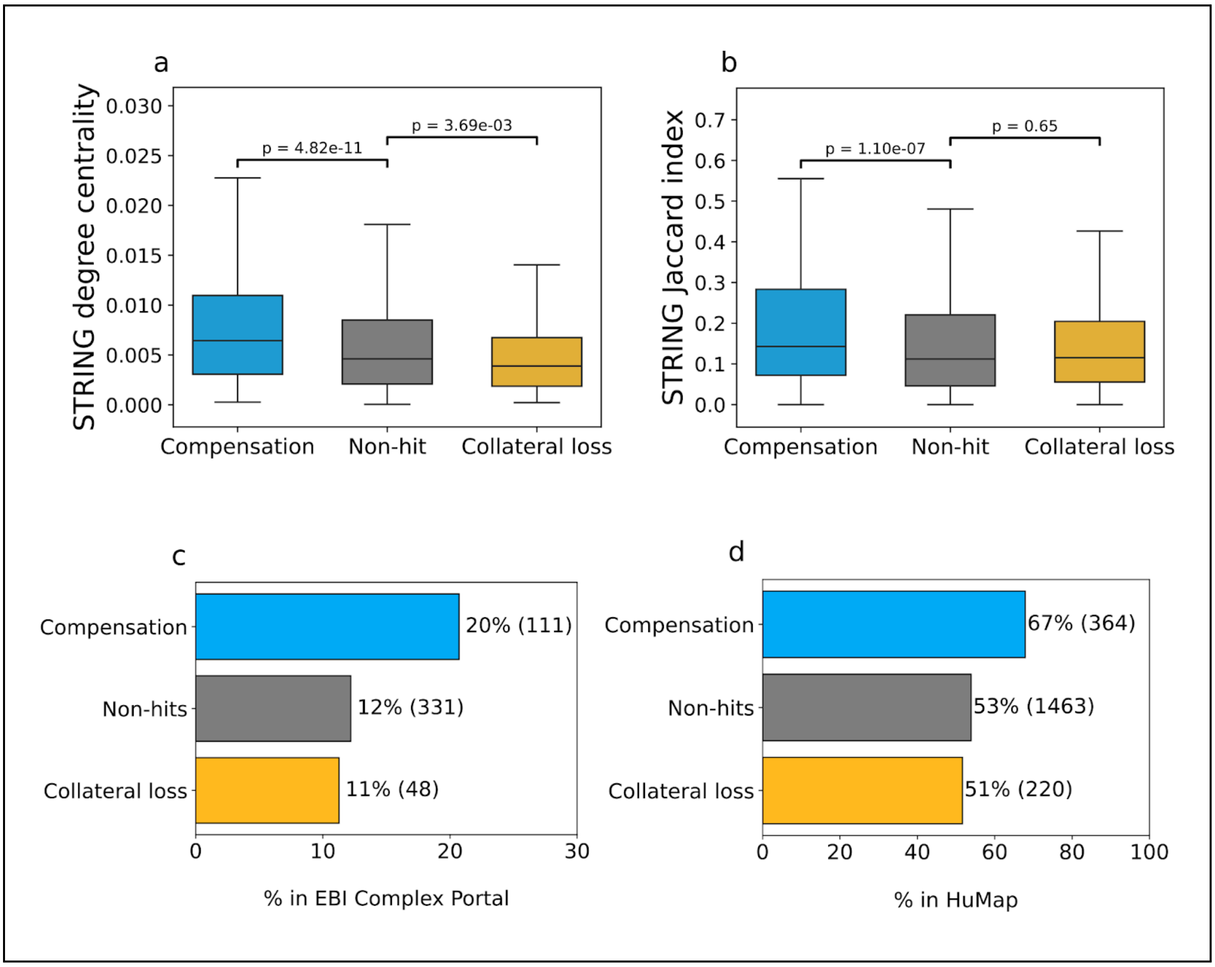


**Supplementary Figure 5. Compensation pairs are more central in the protein-protein interaction network when analyzing the STRING physical subnetwork, EBI ComplexPortal, and the HuMap database. (a)** Boxplot showing distributions of STRING degree centrality for compensation pairs, non-hits, and collateral loss pairs [30]. **(b)** Similar boxplot comparing the distributions of Jaccard indices calculated with the STRING network. **(c)** Barchart comparing the percentage of pairs in each set where either member of the pair is a member of a protein complex in EBI Complex Portal [32]. **(d)** Similar barchart plotted to show the percentages of pairs where either member is in a predicted HuMap complex [33]

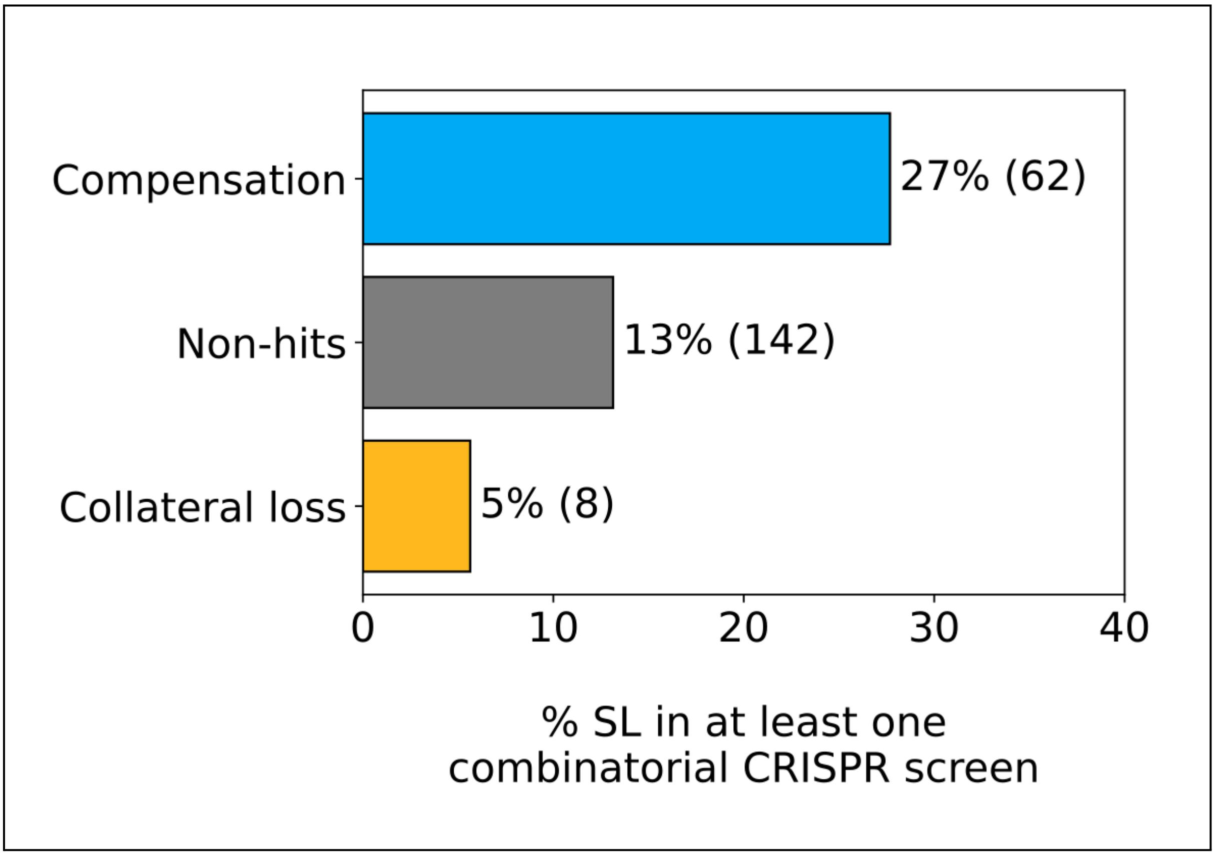


**Supplementary Figure 6. Compensation hits are enriched for paralog pairs that are synthetic lethal in at least one combinatorial CRISPR screen.**

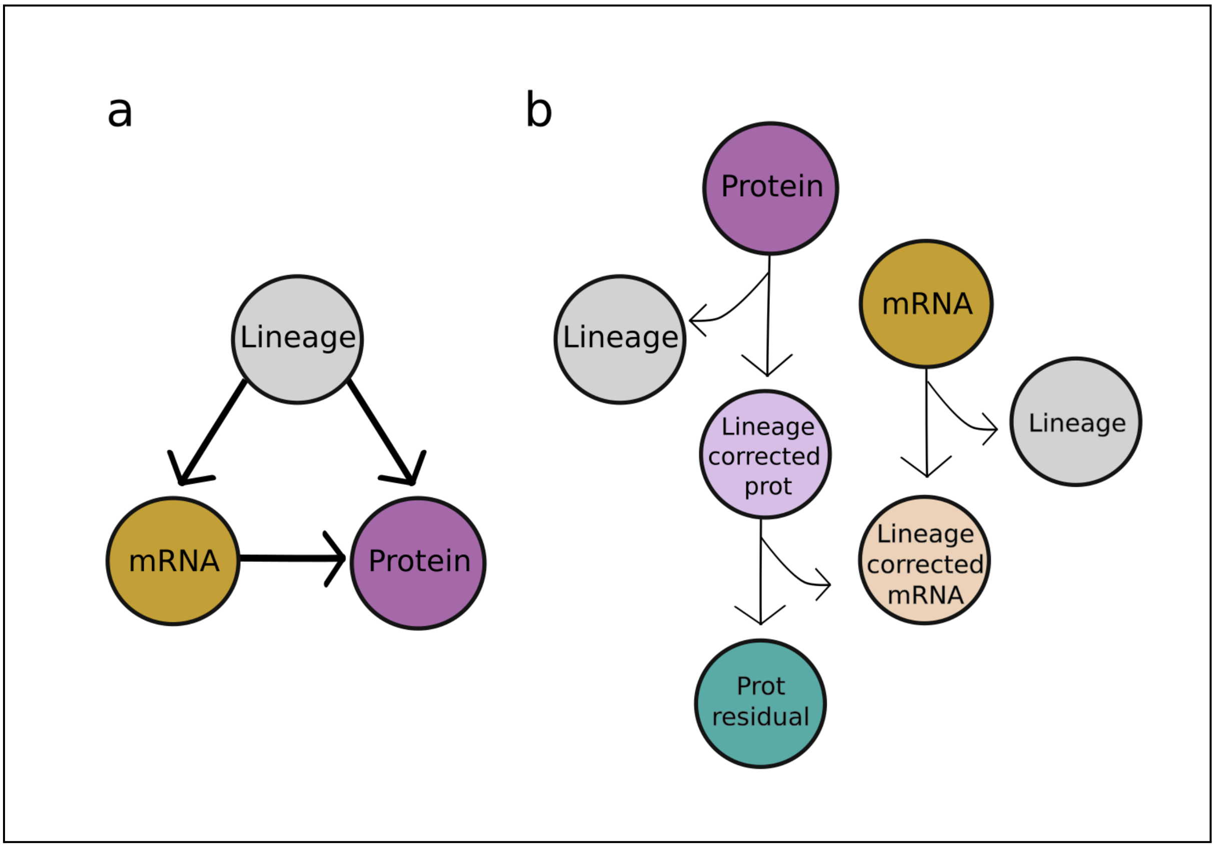


**Supplementary Figure 7. Generation of protein residual dataset by regressing out transcript abundance from protein abundance (a)** Directed Acyclic Graph (DAG) showing the causal links between lineage/study, transcript abundance, and protein abundance. **(b)** Workflow diagram showing how mRNA abundance is regressed out from protein abundance, while taking into account the effect of lineage on both variables.

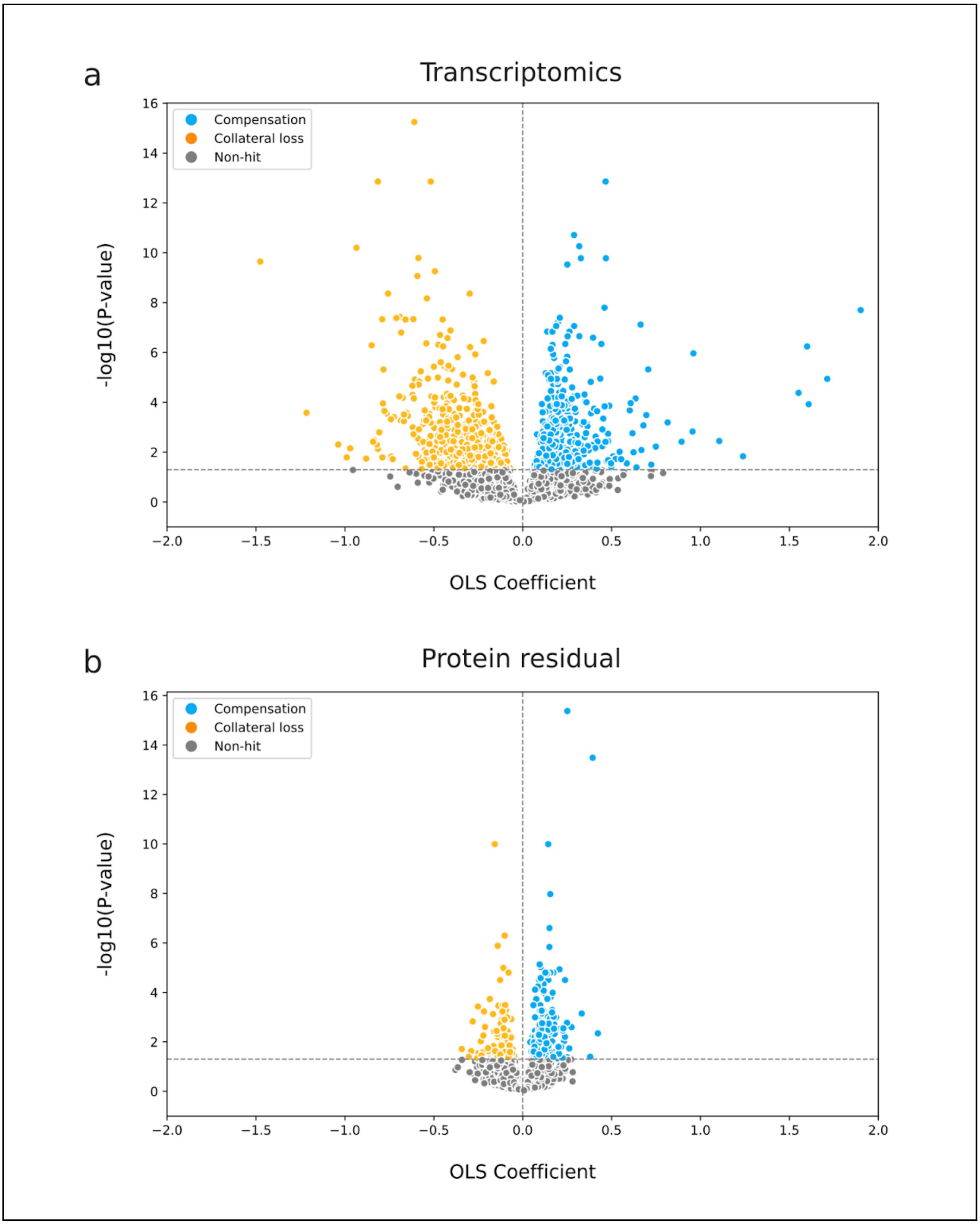


**Supplementary Figure 8. Volcano plots showing results for all paralog tests run with the CPTAC transcriptomic and protein residual datasets.** Volcano plots show -log10(FDR) vs ordinary least squares (OLS) coefficient for all pairs tested. The same 4,568 pairs tested in the proteomic analysis were tested with these datasets.

**Supplementary Table 1:** Processed HAP1 proteomic data.

**Supplementary Table 2:** All self-abundance HAP1 tests.

**Supplementary Table 3:** All paralog HAP1 tests.

**Supplementary Table 4:** All self-abundance CPTAC tests.

**Supplementary Table 5:** CPTAC proteomic results.

**Supplementary Table 6:** CPTAC transcriptomic and protein residual results.

**Supplementary Table 7:** Biological information for all HAP1 pairs.

**Supplementary Table 8:** Biological information for all CPTAC pairs.

**Supplementary Table 9:** All Fishers Exact Test results (categorical overlap tests) for CPTAC pairs.

**Supplementary Table 10:** All t-test results (quantitative overlap tests) for CPTAC pairs.

- All supplementary tables are included in ***paralog_protein_compensation_supptables.xlsx***, with additional information for all tables in **Sheet 1.**

- All references in figures point to articles that have also been referenced in the main text, and so share the same numbering system. No references are unique to figure legends.
